# Plasma membrane abundance dictates phagocytic capacity and functional crosstalk in myeloid cells

**DOI:** 10.1101/2023.09.12.556572

**Authors:** Benjamin Y. Winer, Alexander H. Settle, Alexandrina M. Yakimov, Carlos Jeronimo, Tomi Lazarov, Murray Tipping, Michelle Saoi, Anjelique Sawh, Anna-Liisa L. Sepp, Michael Galiano, Yung Yu Wong, Justin S. A. Perry, Frederic Geissmann, Justin Cross, Ting Zhou, Lance C. Kam, Hilda Amalia Pasoli, Tobias Hohl, Jason G. Cyster, Orion D. Weiner, Morgan Huse

## Abstract

Professional phagocytes like neutrophils and macrophages tightly control what they eat, how much they eat, and when they move after eating. We show that plasma membrane abundance is a key arbiter of these cellular behaviors. Neutrophils and macrophages lacking the G-protein subunit Gβ4 exhibit profound plasma membrane expansion due to enhanced production of sphingolipids. This increased membrane allocation dramatically enhances phagocytosis of bacteria, fungus, apoptotic corpses, and cancer cells. Gβ4 deficient neutrophils are also defective in the normal inhibition of migration following cargo uptake. In Gβ4 knockout mice, myeloid cells exhibit enhanced phagocytosis of inhaled fungal conidia in the lung but also increased trafficking of engulfed pathogens to other organs. These results reveal an unexpected, biophysical control mechanism lying at the heart of myeloid functional decision-making.

## Introduction

Professional phagocytes of the myeloid lineage, including neutrophils and macrophages, maintain homeostasis by clearing apoptotic corpses, cellular debris, and invading pathogens(Wang et al. 2017; Peiseler and Kubes 2019; Witter, Okunnu, and Berg 2016; Pylaeva et al. 2022). These cells are also central components in emerging immunotherapeutic strategies to combat infections, cardiovascular disease, and cancer(Alvey and Discher 2017; Dooling et al. 2023; Andrechak, Dooling, and Discher 2019; Johansson and Kirsebom 2021). Phagocytes take up cargo via phagocytosis, an evolutionarily conserved engulfment process that is triggered by the recognition of cognate cargo ligands, such as phosphatidylserine, complement, or antibodies, by specific receptors on the phagocyte(Gordon 2016). Target recognition elicits a dramatic remodeling of the cytoskeleton, which shapes the overlying plasma membrane into a phagocytic cup that surrounds and then internalizes the cargo(Jaumouille and Wa-terman 2020; Mylvaganam, Freeman, and Grinstein 2021). In carrying out their clearance function, myeloid cells exhibit not only robust cargo uptake but also the ability to coordinate this activity with other cellular behaviors(Bretou et al. 2017; Chabaud et al. 2015; Luo et al. 2009; Branzk et al. 2014; Yipp et al. 2012). Crosstalk between phagocytosis and cell motility is particularly well-established, with studies documenting an antagonistic relationship between cell migration and engulfment responses in multiple myeloid cell types(Bretou et al. 2017; Chabaud et al. 2015; Luo et al. 2009). In neutrophils specifically, transient arrest following phagocytosis is thought to curtail the dissemination of intra-cellular microbes(Jhingran et al. 2012; Hopke et al. 2020; Hamza et al. 2014). Accordingly, any efforts to harness professional phagocytes therapeutically will require a mechanistic understanding of not only phagocytosis itself but also its cross-regulatory effects on other activities.

Most prior research on phagocytosis has focused on the biochemical mechanisms that control it, and as a result, much is now known about the molecules that mediate cargo recognition and the signal transduction pathways that drive cup formation and cargo engulfment(Uribe-Querol and Rosales 2020). Phagocytosis is also an intensely physical process(Vorselen, Labitigan, and Theriot 2020; Jaumouille and Waterman 2020), implying that it might be subject to biophysical as well as biochemical modes of regulation. In that regard, it is interesting to note that the deformability of both cargo and underlying substrate have been shown to modulate the engulfment behavior of macrophages(Jain, Moeller, and Vogel 2019; Adlerz, Aranda-Espinoza, and Hayenga 2016; Blakney, Swartzlander, and Bryant 2012; Okamoto et al. 2018; Underhill and Goodridge 2012; Sosale et al. 2015). Whether the architecture and mechanics of the phagocyte itself might also influence cargo uptake is not known, however, and how cell-intrinsic properties of this kind might affect functional crosstalk between phagocytosis and other cel-lular behaviors is completely unexplored.

## Results

### G*β*4 deficiency enhances phagocytic responses against a wide range of targets

To identify novel mechanisms controlling phagocytosis and functional crosstalk, we performed a candidate screen focused on heterotrimeric G-protein signaling, which regulates phagocytosis and migration in multiple myeloid cell types(Peracino et al. 1998; Wen et al. 2019; Pan et al. 2016; Cohen et al. 2022; Kamber et al. 2021; Yamada and Sixt 2019; Schwarz et al. 2017; Sarris and Sixt 2015). Activating signals induce the dissociation of the G-protein *α* subunit (G*α*) from the *βγ* subcomplex (G*βγ*), freeing both components to bind and activate specific downstream effector molecules(Clapham and Neer 1997). The bulk of prior research in this area has focused on G*α* isoforms, while comparatively less is known about specific G*β* and G*γ* subunits(Kankanamge et al. 2022). Accordingly, we applied CRISPR-Cas9 to knock out each G*β* subunit in human neutrophil-like HL-60 cells (Fig. 1a and fig. S1a and b). Gene targeting was carried out in self-renewing HL-60 precursors, which were then differentiated into neutrophil-like cells by the addition of DMSO. All terminally differentiated cells were CD15+ CD16+ (fig. S1c), indicating that the genetic modifications we introduced did not prevent HL-60 progenitors from becoming neutrophils.

**Fig. 1.**
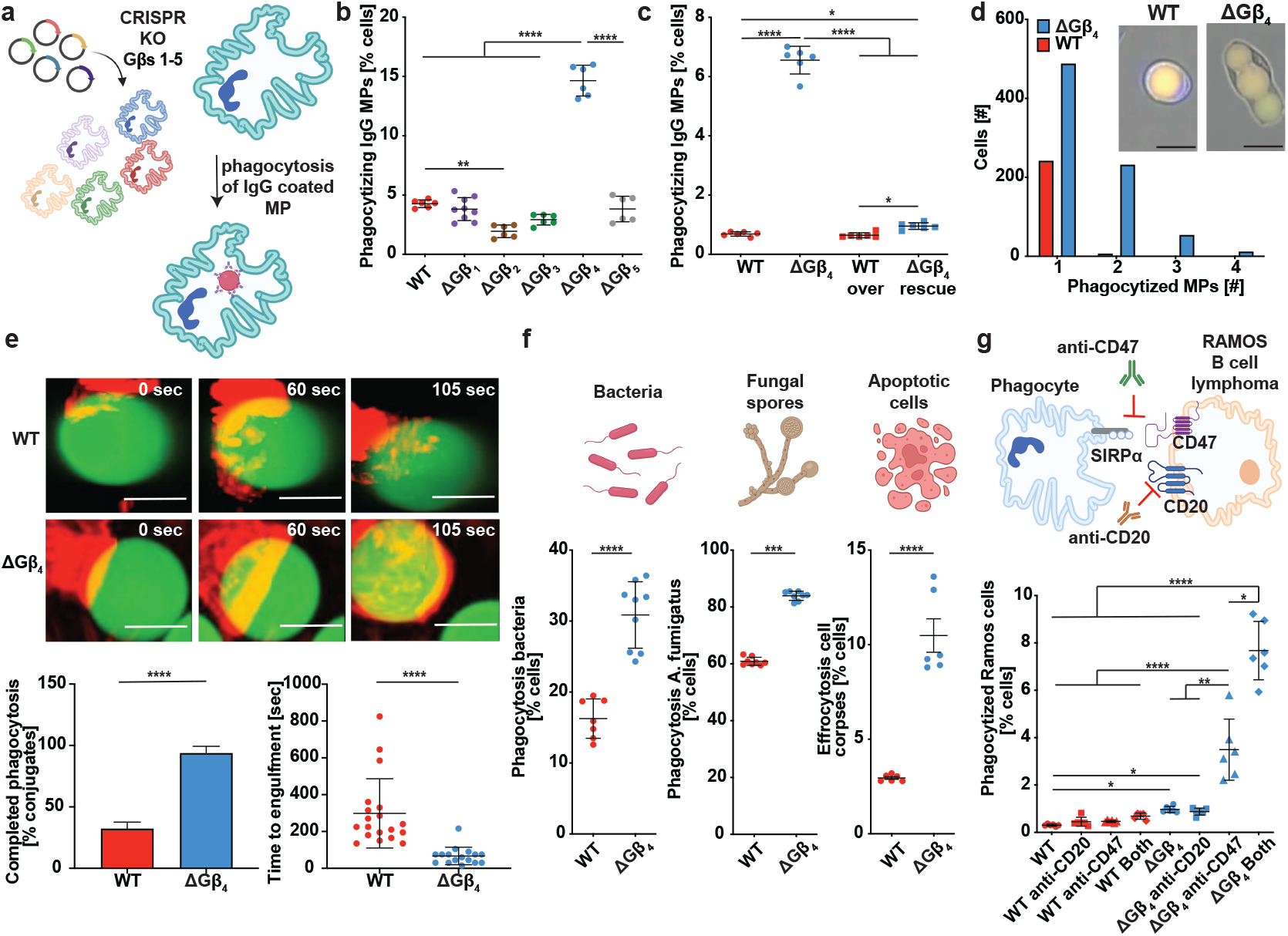
ΔGβ4 neutrophils have increased phagocytic capacity for a wide range of targets. a, Schematic of Gβ knock-out workflow in HL-60 cells and the MP-based phagocytosis assay. b, WT and ΔGβ HL-60 cells were challenged with human IgG coated MPs and phagocytosis quantified after 3 h. Data are mean s.d. of three experiments. One-way ANOVA, ** p< 0.01, ****p< 0.0001. c, WT and ΔGΔ4 HL-60 cells transfected with exogenous Gβ4 (WT over and DGb4 rescue) or control lentivirus (WT and DGb4) were challenged with human IgG coated MPs and phagocytosis quantified after 3 h. Data are mean s.d. of three experiments. One-way ANOVA, * p< 0.05 and ****p< 0.0001. d, ΔGβ4 and WT HL-60 cells were challenged with IgA coated MPs and imaged over 3 hours to assess per cell phagocytic capacity. Histogram shows the number of ΔGβ4 and WT cells that consumed 1, 2, 3, or 4 MPs during the experiment. Inset shows sample images of WT and ΔGβ4 cells after MP uptake. Scale bars = 20 μm. e, WT and ΔGβ4 HL-60 cells expressing F-tractin (red) were fed IgG coated MPs (green) and phagocytic uptake was monitored for 10-30 minutes. Above, representative interactions are shown in time-lapse montage format. Scale bars = 5 μm. Below, quantification of the frequency (left) and speed (right) of phagocytosis. Data are mean s.d. of 6 experiments. Unpaired t-test, ****p< 0.0001. f, βGΔ4 and WT HL-60 cells were challenged with fluorescently labeled Pseudomonas aeruginosa (left), Aspergillus fumigatus (center), or apoptotic Jurkat cell corpses (right), and phagocytosis quantified after 3 h. Data are the mean s.d. from 3 separate experiments. Unpaired t-test *** p< 0.001, **** p< 0.0001. g, Above, schematic of anti-CD47 and anti-CD20 induced uptake of RAMOS B lymphoma cells by phagocytes. Below, DGb4 and WT HL-60 cells were challenged with RAMOS B lymphoma cells in the presence or absence of anti-CD20 and anti-CD47 as indicated. Data are mean s.d. of 3 experiments. One-way ANOVA, * p< 0.05, ** p< 0.01, **** p< 0.0001.

Knockout and control HL-60 cells were then subjected to phagocytosis assays. As cargo for these experiments, we prepared 10 μm diameter IgG-coated polyacrylamide microparticles (MPs)(Vorselen et al. 2020; Vorselen et al. 2021) derivatized with the fluorescent dyes FITC and LRB (Fig. 1a and fig. S2). FITC, but not LRB, fluorescence is quenched in acidic phagolysosomes, producing a color change that can be monitored by flow cytometry and fluorescence microscopy. In this manner, we were able to identify G*β*subunits that either positively or negatively regulated phagocytosis. Consistent with prior work(Cohen et al. 2022; Kamber et al. 2021), HL-60 cells lacking G*β*2 exhibited a 50 percent reduction in particle uptake (Fig. 1b). By contrast, depletion of G*β*4 induced a striking 3- to 4-fold increase in phagocytosis relative to wild type (WT) controls (Fig. 1b). This hyperphagic behavior was readily apparent in both monocultures as well as cocultures containing a 1:1 mixture of WT and G*β*4 knockout (ΔG*β*4) HL-60 cells (fig. S3a-b), indicating that the phenotype was cell intrinsic. To ensure that our CRISPR-editing did not generate any off-target effects, we verified that re-expression of G*β*4 on the knock-out background restored phagocytosis to WT levels (Fig. 1c, fig. S3c). Using live imaging, we found that ΔG*β*4 deficiency markedly enhanced particle consumption on a per-cell basis; while the vast majority of phagocytic WT cells took up just one MP, almost half of phagocytic. ΔG*β*4 cells engulfed two or more MPs (Fig. 1d). We next leveraged super-resolution microscopy to examine engulfment dynamics in more detail. ΔG*β*4 HL-60 cells formed stereotypical phagocytic cups featuring a pro-nounced band of filamentous actin (F-actin) at the leading edge (Fig. 1e). This morphology was much less apparent in WT cells, which still engaged MP targets but only achieved full engulfment in less than 50 percent of conjugates. In contrast, ΔG*β*4 HL-60 cells completed over 90 percent of their phagocytic attempts and executed engulfment more than twice as fast as their wild-type counterparts (Fig. 1e, Supplementary Video 1).

To further explore the scope of the Δ.G*β*4 phenotype, we challenged WT and ΔG*β*4 HL-60 cells with MPs bearing alternative coatings, namely phosphatidylserine (PS), complement, IgG, and IgA. ΔG*β*4 cells phagocytosed 2-to 10-fold more cargo in each case, indicating that their hyperphagy was not limited to a specific uptake receptor (fig. S3d). Next, we measured the phagocytosis of four distinct biological targets: the gram-negative bacterium Pseudomonas aeruginosa, Staphylococcus aureus bioparticles, conidia from the fungus Aspergillus fumigatus, and apoptotic Jurkat T cell corpses (Fig. 1f, fig. S3e-f). ΔG*β*4 HL-60 cells significantly outperformed controls in every case, further supporting the idea that G*β*4 deficiency potentiates phagocytosis against diverse biological cargos. We next investigated whether loss of G*β*4 augments the therapeutic phagocytosis of tumors. Many cancer cells evade immune detection and clearance by expressing CD47, a cell surface protein that functions as a “don’t eat me” signal(Chao, Weissman, and Majeti 2012; Logtenberg, Scheeren, and Schumacher 2020). In certain cases, inhibiting this “don’t eat me” signal with blocking antibodies or peptides against CD47 or its receptor, SIRP*α*, is sufficient to stimulate an engulfment response by phagocytes (Fig. 1g) (Willingham et al. 2012; Jalil et al. 2020; Rodriguez et al. 2013). To adapt this approach to our experimental system, we challenged HL-60 cells with Ramos B lymphoma target cells in the absence or presence of anti-CD47 blockade. In some samples, opsonizing antibodies against the B cell marker CD20 were added to further promote phagocytosis via Fc receptor engagement. WT HL-60 cells did not take up Ramos cells, and treatment with anti-CD47 and/or anti-CD20 failed to enhance their activity. In contrast, G*β*4 HL-60 cells exhibited a low, but measurable level of baseline phagocytosis, which increased sharply (3-fold) in the presence of anti-CD47, and even more so (8-fold) when anti-CD47 was combined with anti-CD20 (Fig. 1g). Collectively, these results demonstrate that G4 deficiency substantially augments phagocytic responses across a wide range of targets.

### G*β*4 deficiency alters cell migration and crosstalk between phagocytosis and motility

As proper coordination between cargo uptake and motility is essential for myeloid cell function(Bretou et al. 2017; Chabaud et al. 2015; Luo et al. 2009), we next investigated the effects of G*β*4 on migratory capacity. To this end, we employed a chemotaxis assay in which HL-60 cells were attached to fibronectin-coated glass and then presented with a point source of the peptide chemoattractant fMLF, applied via micropipette (Fig. 2a). Under these conditions, neutrophils adopt a fan-like morphology with a broad leading edge directed toward the point source and a thin uropod trail-ing behind. Both ΔG*β*4 and WT cells were able to migrate up the fMLF gradient, but ΔG*β*4 cells did so at a significantly slower rate (Fig. 2b, Supplementary Video 2). This reduction in motility was associated with the formation of extended “tails” at the rear of migrating ΔG*β*4 cells, suggestive of a defect in uropod retraction (Fig. 2c). Indeed, we observed multiple instances in which extensively elongated ΔG*β*4 cells appeared to be “struggling” against their own uropod to make forward progress (Fig. 2c). Next, we in-terrogated crosstalk between phagocytosis and migration by quantifying the motility of HL-60 cells after uptake of one IgG-coated MP. WT and ΔG*β*4 neutrophils were fed MPs with distinct fluorescent labels (pHrodoRed (WT) versus pHrodoGreen (ΔG*β*4)) to facilitate imaging and tracking in mixed samples. Particle consumption slowed the movement of WT neutrophils considerably relative to unfed, WT controls (Fig. 2b). The same could not be said for ΔG*β*4 neutrophils, which moved faster than their unfed ΔG*β*4 counterparts (Fig. 2b). Although the instantaneous speed of fed ΔG*β*4 neutrophils remained slightly less than that of fed WT cells, their migration was more persistent, yielding signifi-cantly longer tracks that more closely approached the fMLF point source (Fig. 2d-e, Supplementary Video 2). These motility phenotypes were mirrored by changes in cell shape. In WT neutrophils, phagocytosis appeared to hamper migratory polarization; cells periodically collapsed into a rounded configuration, and these morphological changes tended to co-incide with stalls in motility (Fig. 2c, Supplementary Video 2). By contrast, ΔG*β*4 morphology was normalized by particle uptake; fed cells formed persistent leading edges and lacked the extended uropods characteristic of the unfed state (Fig. 2c). Hence, ΔG*β*4 depletion alters not only the speed of neutrophil migration but also the ability of phagocytosis to inhibit motility.

**Fig. 2.**
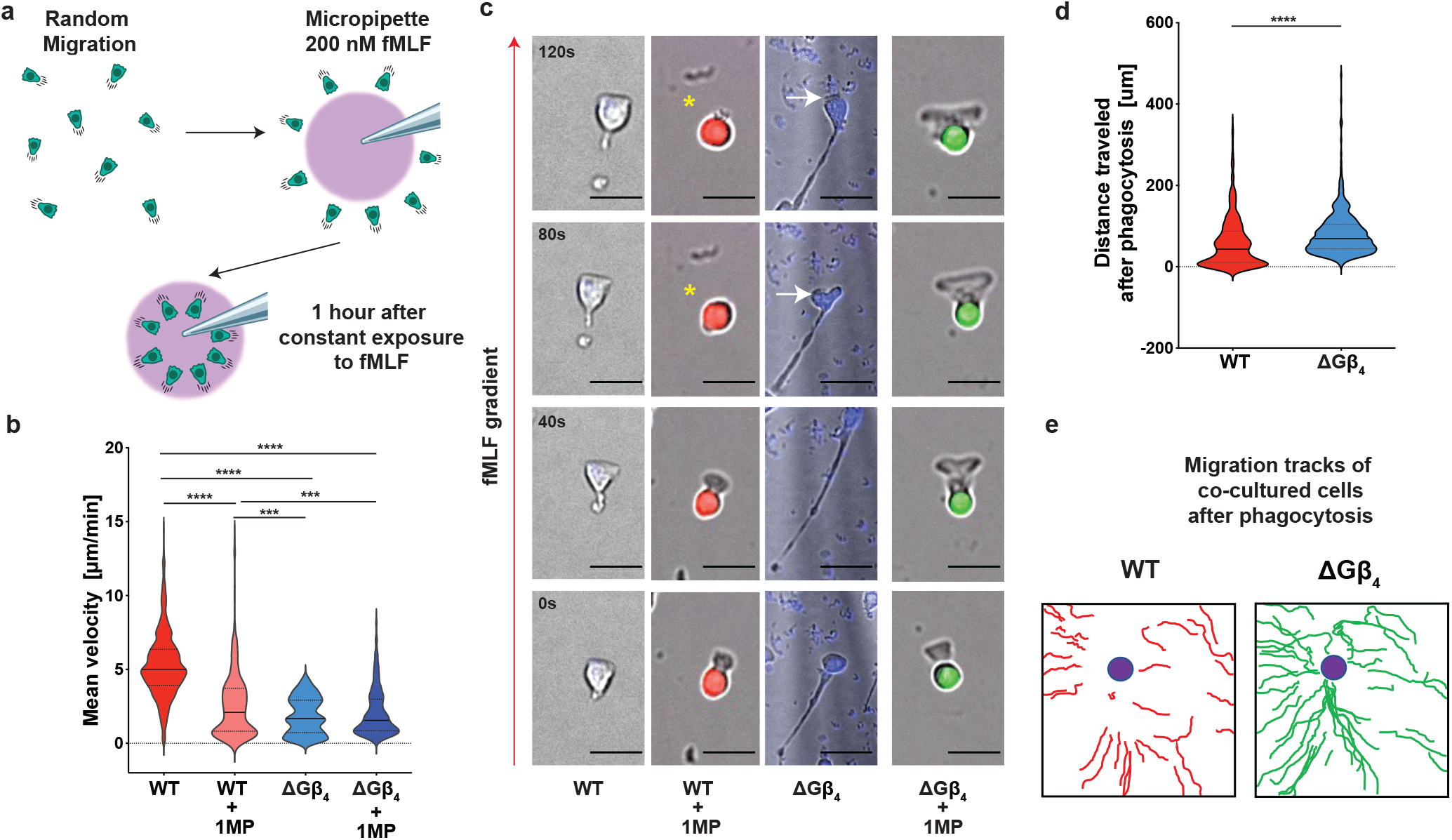
Gβ4 deficiency alters cell migration and crosstalk between phagocytosis and motility. a. Schematic of the micropipette-based chemotaxis assay. b-d, ΔGβ4 and WT HL-60 cells were seeded onto a fibronectin coated slide and exposed to 200 nM fMLF delivered through a micropipette over the course of an hour. Alternatively, ΔGβ4 and WT cells that had consumed 1 MP were sorted, seeded at a 1:1 ratio on fibronectin coated glass, and exposed to 200 nM fMLF through a micropipette for one hour. b, Mean velocity of WT, WT + 1MP, ΔGβ4, and ΔGβ4 + 1MP cells was calculated from 3 separate 1 hour videos. One-way ANOVA, *** p< 0.001, and **** p< 0.0001. c, Time-lapse montages of representative cells in each experimental group, with the red arrow indicating the direction of the fMLF gradient. Yellow asterisks denote a transient loss of migratory cell polarity in a WT + 1MP cell. White arrows indicate cell body regression in a ΔGβ4 cell with an extended uropod. Scale bars = 20 μm. d, Quantification of total distance travelled by WT + 1MP and ΔGβ4 + 1MP cells after 1 hour of exposure to 200 nM fMLF. Unpaired t test **** p< 0.0001. e, Representative migration tracks for WT + 1MP and ΔGβ4 + 1MP over the course of 1 hour exposure to 200 nM fMLF. Violins in b and d encompass the entire distribution, with solid horizontal lines indicating the median and dotted lines indicating the upper and lower quartiles.

### G*β*4 deficiency alters lipid composition and plasma membrane abundance

The striking uropod extension displayed by migrating ΔG*β*4 HL-60 cells was suggestive of a substantial change in their cellular architecture. Consistent with this notion, scanning electron microscopy (SEM) indicated that ΔG*β*4 HL-60 cells had a more ruffled surface appearance than WT controls (Fig. 3a). To investigate this structural difference more closely, we performed Focused Ion Beam (FIB)-SEM, a method in which successive sections from the same sample are imaged by SEM and then used to generate a nanometer-resolution 3-D reconstruction(Ritter et al. 2022). WT and ΔG*β*4 HL-60 cells were osmium-stained to highlight lipid-rich cellular components, embedded in resin, and subjected to FIB-SEM at 40 nm sectioning. Supervised machine learning was used to define the plasma membrane in each image, followed by 3-D rendering(Wang et al. 2023). The resulting reconstructions revealed striking differences in plasma membrane configuration. Whereas the surface of WT cells was mostly smooth with small extensions, ΔG*β*4 cells exhibited large flaps of plasma membrane projecting up to 10 μm from the cell body (Fig. 3b, Supplementary Video 3). The presence of these structures increased the surface to volume ratio of ΔG*β*4 HL-60 cells by a factor of two relative to WT controls (Fig. 3c). ΔG*β*4 cells also contained larger and more numerous cytoplasmic lipid droplets, which appeared as high contrast compartments in osmium-stained FIB-SEM images (Fig. 3b, 3d-e). This increase in lipid droplet load was confirmed by BODIPY staining (fig. S4a-b). Hence, *β*4 depletion increases plasma membrane abundance and promotes lipid droplet formation in HL-60 cells.

**Fig. 3.**
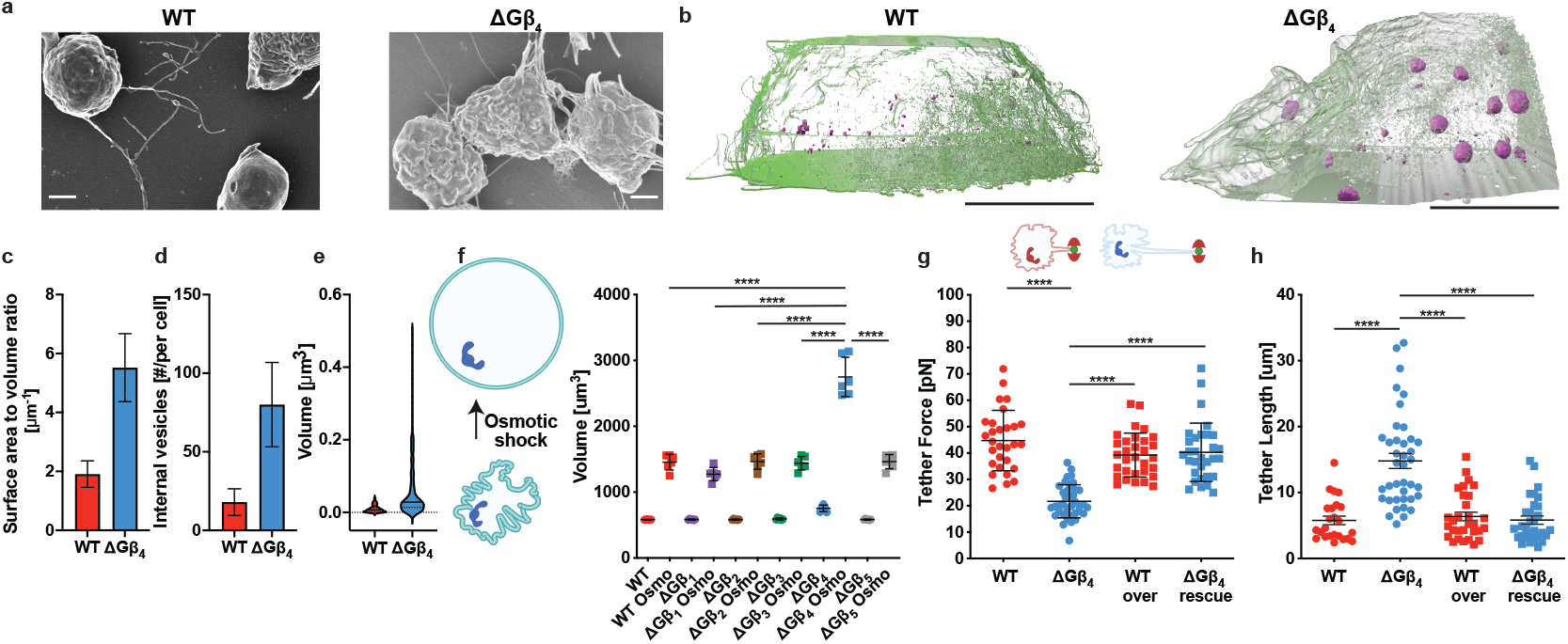
ΔGβ4 HL-60 cells have increased plasma membrane and decreased membrane tension. a, Representative SEM images of WT and βGΔ4 HL-60 cells. Scale bars = 2 μm. b-e, βGΔ4 and WT HL-60 cells stained with potassium ferrocyanide and osmium tetroxide were imaged by FIB-SEM (n = 2 of each cell type). b, 3-D reconstructions of representative cells, with plasma membrane shown in semi-transparent green and lipid droplets in solid magenta. Scale bars = 5 μm. c, Quantification of surface area to volume ratio. d, Quantification of lipid droplet number per cell. In c and d, data are mean s.d. e, Quantification of lipid droplet volume (n = 20 droplets in WT cells and 24 droplets in βGΔ4 cells). Violins encompass the entire distribution, with solid horizontal lines indicating the median and dotted lines indicating the upper and lower quartiles. f, HL-60 cells lacking the indicated Gb subunits, along with WT controls, were subjected to osmotic shock to induce swelling to maximal volume. Left, schematic of the approach. Right, volume measurements before and after osmotic shock (Osmo). Data are mean s.e.m. of three experiments. One-way ANOVA ****p< 0.0001. g-h, Membrane tethers were generated from WT and ΔGβ4 HL-60 cells transduced with exogenous Gβ4 (WT over and ΔGβ4 rescue) or control lentivirus (WT and ΔGβ4). g, Above, schematic of the tether pulling process. Below, quantification of membrane tension. h, Quantification of membrane tether length. Data in g and h are mean s.d. of five experiments. One-way ANOVA, ****p< 0.0001. WT (n=29), ΔGβ4 (n=39), βGΔ4 rescue (n=31), and WT over (n=31).

To better quantify the ΔG*β*4 membrane accumulation phenotype, we exposed WT and ΔG*β*4 HL-60 cells to osmotic shock, which swells cells to the limits of their plasma membrane capacity and thereby enables comparisons of total surface area. ΔG*β*4 HL-60 cells were only slightly larger than their WT counterparts under isotonic conditions. Upon transfer to hypotonic medium, however, ΔG*β*4 cells expanded to twice the size of controls (Fig. 3f). Assuming that swelled HL-60 cells are spherical, this volume differential implies a 60 percent increase in plasma membrane surface area. This excess plasma membrane would be expected to facilitate the formation of phagocytic cups and thereby promote engulfment. To quantify membrane mobilization during cup formation, we utilized a “frustrated phagocytosis” assay(Masters et al. 2013; Kovari et al. 2016) in which HL-60 cells were applied to IgG-coated glass slides. Neutrophils and macrophages form flat, unresolved phagocytic cups on surfaces of this kind, which are easily visualized by total internal reflection fluorescence (TIRF) microscopy. Frustrated phagosomes formed by ΔG*β*4 HL-60 cells were substantially larger than those of WT controls (fig. S4c), consistent with a role for increased membrane abundance in phagocytic cup formation.

Membrane tension is known to inhibit both phagocytosis and migration, presumably by mechanically constraining the formation of actin-based structures and/or activating inhibitory mechanosensory pathways(Masters et al. 2013). We speculated that, by increasing plasma membrane abundance, *β*4 deficiency might reduce membrane tension and thereby attenuate these regulatory effects. To explore this possibility, we utilized an established approach in which a concanavalin-A-coated bead is adsorbed to the cell surface and then pulled away using an optical trap(Dai and Sheetz 1995). The displaced bead remains attached to the cell via a thin membrane tether, which exerts a restoring force that is proportional to the square of membrane tension(Bo and Waugh 1989; Heinrich and Waugh 1996). In this manner, we found that *β*4 deficiency reduced tether forces by a factor of two (Fig. 3g). ΔG*β*4 cells also generated longer tethers then WT controls, indicating that they possessed larger plasma membrane reservoirs (Fig. 3h)(Raucher and Sheetz 1999). Rescue experiments confirmed that both phenotypes were specific to *β*4 (Fig. 3g-h). In conjunction with the imaging and osmotic shock studies described above, these results suggest a central role for plasma membrane abundance in the regulation of neutrophil effector responses.

To explore potential mechanisms underlying the ΔG*β*4 plasma membrane phenotype, we subjected ΔG*β*4 and WT HL-60 cells to comparative lipidomics (Fig. 4a and fig. S5a-b). As expected, ΔG*β*4 HL-60 cells contained more total lipids than WT controls on a per cell basis (fig. S5c). This increase did not apply to all lipid subtypes, however, as many species were unchanged between samples and some, like hexosylceramides, were more abundant in WT cells (fig. S5d). ΔG*β*4 neutrophils were distinguished by disproportionate enrichment of sphingolipids, particularly ceramide and sphingomyelin (Fig. 4a and fig. S5c), which are predominantly found in the plasma membrane. Using RNA-sequencing, we documented selective upregulation of sphingolipid synthesis genes in ΔG*β*4 neutrophils after MP feeding (Fig. 4b), establishing an additional link between sphingolipids and the ΔG*β*4 phenotype. To interrogate the role of sphingolipids in plasma membrane expansion more directly, we asked whether inhibition of sphingolipid synthesis could reverse the effects of *β*4 deficiency. To this end, we applied the small molecule myriocin, a potent inhibitor of serine palmitoyl transferase, the first enzyme in the sphingolipid synthesis pathway (Fig. 4c and fig. S5e)(Glaros et al. 2007; Wadsworth et al. 2013). Myriocin treatment during differentiation restored the membrane tension of ΔG*β*4 HL-60 cells to WT levels (Fig. 4d), and it also largely reversed their hyperphagic behavior (Fig. 4e). Notably, acute treatment of fully differentiated ΔG*β*4 HL-60 cells with myriocin had no effect on MP uptake (fig. S5f), implying that the functionally relevant effects of *β*4 signaling on sphingolipid synthesis occur during the differentiation phase. Collectively, these results indicate that the excess plasma membrane characteristic of ΔG*β*4 cells is due, at least in part, to increased sphingolipid synthesis.

**Fig. 4.**
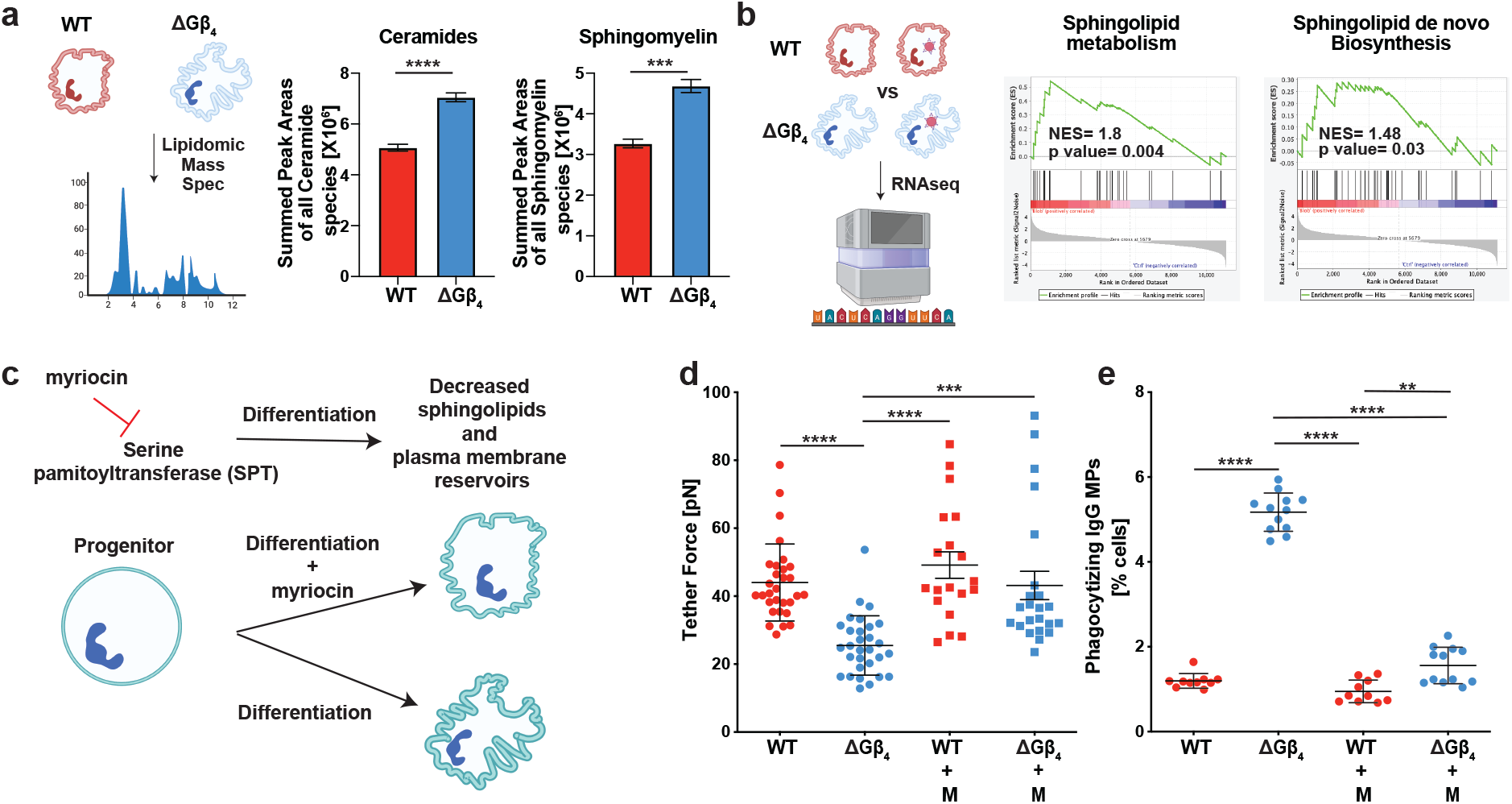
Gβ4 regulates plasma membrane expansion via sphingolipid synthesis. a, Left, schematic overview of the lipidomics workflow. Graphs show quantification of ceramides (center) and sphingomyelin (right). Data are mean s.d. of three biological replicates. Unpaired t-test, ***p< 0.001, ****p< 0.0001. b, Left, schematic overview of RNA-seq workflow. ΔGβ4 and WT cells were compared before and after consumption of IgG coated MPs. Center and right, gene set enrichment analyses associating the gene expression changes induced by MP uptake in βGΔ4 cells with gene sets for sphingolipid metabolism (center) and sphingolipid de novo biosynthesis. NES = normalized enrichment score. c, Schematic showing the anticipated effects of myriocin treatment on sphingolipid synthesis and plasma membrane abundance. d-e, WT and βGΔ4 HL-60 neutrophils were differentiated in the presence or absence of myriocin (M), and then subjected to biophysical and functional analysis. d, Quantification of membrane tension, determined by membrane tether pulling. Data represent mean s.d. of 5 experiments. One-way ANOVA, *** p< 0.001 and **** p< 0.0001. e, Cells were challenged with IgG coated MPs and phagocytosis quantified after 3 h. Data represent mean s.d. of 3 experiments. One-way ANOVA, *** p< 0.001 and **** p< 0.0001.

### G*β*4 controls membrane abundance and phagocytosis in primary myeloid cells

Having demonstrated ΔG*β*4-induced hyperphagy in HL-60 cells, we next investigated whether this mechanism oper-ates in primary neutrophils and macrophages. By applying CRISPR-Cas9 targeting to the ER-HoxB8 immortalized murine neutrophil progenitor system(Hopke et al. 2020; Greene et al. 2022; Khoyratty et al. 2021) (fig. S6a-b), we were able to generate primary neutrophils lacking *β*4, along with controls expressing non-targeting guide (g)RNA. Both ΔG*β*4 and WT ER-HoxB8 neutrophils expressed equivalent levels of CD11b and Ly6G after differentiation (fig. S6c), indicating that Gb4 is dispensable for the acquisition of neutrophil cell fate. As with ΔG*β*4 HL-60 cells, ΔG*β*4 ER-HoxB8 neutrophils displayed clear signs of plasma membrane dysregulation, including reduced membrane tension and increased membrane tether length in optical trap experiments (fig. S6d-e). ΔG*β*4 ER-HoxB8 neutrophils also expanded to nearly twice the volume of WT controls upon transfer to hypotonic medium (fig. S6f). In phagocytosis assays, both WT and ΔG*β*4 ER-HoxB8 neutrophils failed to consume IgG-coated MPs, likely because murine neutrophils are substantially smaller than HL-60 cells and therefore unable to accommodate larger (10 μm diameter) cargos. When challenged with smaller (2 μm diameter) S. aureus bioparticles, however, ΔG*β*4 ER-HoxB8 neutrophils more than doubled the consumption of WT controls (fig. S6g).

To assess the role of *β*4 signaling in primary macrophages, we prepared WT and ΔG*β*4 macrophages from human induced pluripotent stem cells (hiPSCs) (Fig. 5a and fig. S7a)(Zhang and Reilly 2017; Vaughan-Jackson et al. 2021; Alsinet et al. 2022). CRISPR-Cas9 was used to target the GNB4 locus (encoding*β*4) at the hiPSC stage, and WT control hiPSCs were generated in parallel using a nontargeting gRNA (fig. S7b). Granulocyte monocyte progenitor (GMP) cells derived from these hiPSC lines were then differ-entiated into mature CD11b+ CD14+ CD68+ macrophages. Importantly, loss of *β*4 did not affect the efficacy of this differentiation protocol (fig. S8). In optical trap experiments, ΔG*β*4 macrophages generated weaker tether forces and longer membrane tethers than WT controls, indicative of reduced membrane tension and larger membrane reservoirs, respectively (Fig. 5b-c). ΔG*β*4 macrophages also became substantially larger than their WT counterparts in hypotonic medium, despite being similarly sized under isotonic conditions (Fig. 5d). These data, which mirrored our results in the HL-60 and ER-HoxB8 systems, strongly suggested that ΔG*β*4 macrophages possess excess plasma membrane. Finally, we evaluated phagocytic capacity and found that *β*4 deficiency markedly increased the uptake of IgG-coated MPs (Fig. 5e). We conclude that the *β*4 pathway regulates plasma membrane abundance and phagocytosis in primary neutrophils and macrophages.

**Fig. 5.**
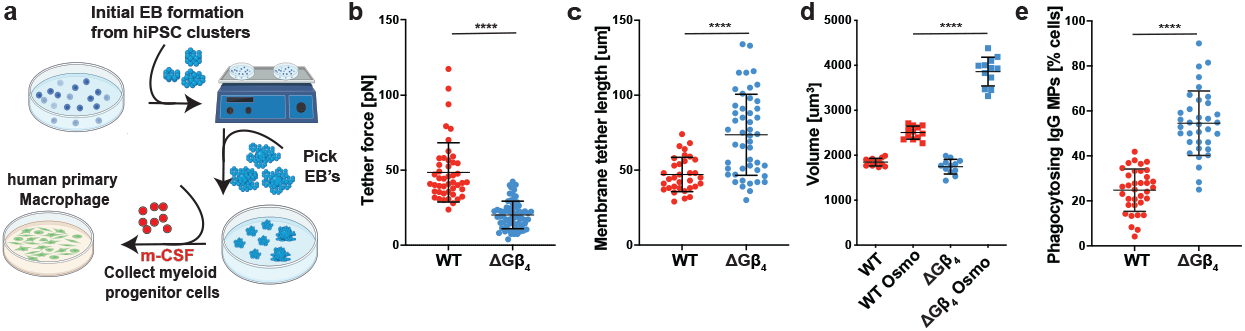
Gβ4 deficiency boosts primary macrophage phagocytosis. a-d, Isogenic ΔGβ4 and WT macrophages were derived from hiPSCs and subjected to biophysical and functional assays. a, Schematic of embryoid body generation from hiPSCs and differentiation of primary macrophages from GMPs. b-c, Membrane tethers were generated from WT (n=46) and DGb4 (n=64) macrophages using an optical trap. b, Quantification of membrane tether force. c, Quantification of membrane tether length. Data in b-c represent mean s.d. of 5 experiments. Unpaired t test **** p< 0.0001. d, ΔGΔ4 and WT macrophages were osmotically shocked and allowed to expand to their full volume to assess total plasma membrane. Data is from 3 separate experiments with one-way ANOVA analysis, **** p< 0.0001. e, WT and ΔGΔ4 macrophages were challenged with IgG coated MPs for 2 h and phagocytic uptake quantified by florescence microscopy. Data represent mean s.d. of 5 experiments. Unpaired t test **** p< 0.0001.

### G*β*4 regulates anti-fungal immunity in vivo

Phagocytes are the first line of defense against microbes in multiple epithelial tissues(Witter, Okunnu, and Berg 2016; Johansson and Kirsebom 2021; Giese, Hind, and Huttenlocher 2019). To evaluate the role of Gβ4-dependent membrane allocation during this early phase of immunity, we generated mice with a targeted deletion of exon 4 of the Gnb4 locus (fig. S9a-b). This modification prematurely terminated the Gn β 4 open reading frame, leading to constitutive Gb4 deficiency. Mice homozygous for this deletion (Gnb4-/-) and WT (Gnb4+/+) littermate controls were infected intratracheally with Aspergillus fumigatus, a fungal pathogen that elicits the robust recruitment of phagocytically active neutrophils to the lung(Guo et al. 2020; Mircescu et al. 2009; Espinosa et al. 2022). To enable flow cytometric detection of fungal uptake by these cells, we employed Fluorescent Aspergillus Reporter (FLARE) conidia that expressed dsRed, a degradable fluorescent protein, and were also labeled with Alexa Fluor (AF)633, a non-degradable small molecule dye. Cells that take up FLARE conidia turn AF633+dsRed+, while cells that have killed the phagocytized fungi become AF633+dsRed-(Fig. 6a)(Jhingran et al. 2012). A. fumigatus infection induced similar levels of neutrophil recruitment to the lungs of WT and Gnb4-/- animals, with CD11b+Ly6G+SiglecF-cells accounting for 70 percent of CD45+ infiltrates, in both experimental groups (Fig. 6b). Fungal phagocytosis was almost 3 times higher in Gnb4-/- neutrophils (Fig. 6c), however, mirroring the hyperphagic phenotype seen in HL-60 cells, ER-HoxB8 neutrophils, and hiPSC derived macrophages lacking Gβ4. Among these phagocytic cells, the frequency of fungal killing was unaffected by Gβ4 deficiency (fig. S9c), consistent with the inter-pretation that Gβ4 signaling modulates cargo uptake and not subsequent acidification of the phagolysosome. Gnb4-/- animals also exhibited a 10 fold reduction in fungal CFU in the lung relative to WT controls at the 18 hour time point (Fig. 6d), indicating that the increased conidial phagocytosis conferred by loss of Gβ4 improved fungal clearance during early stage infection.

**Fig. 6.**
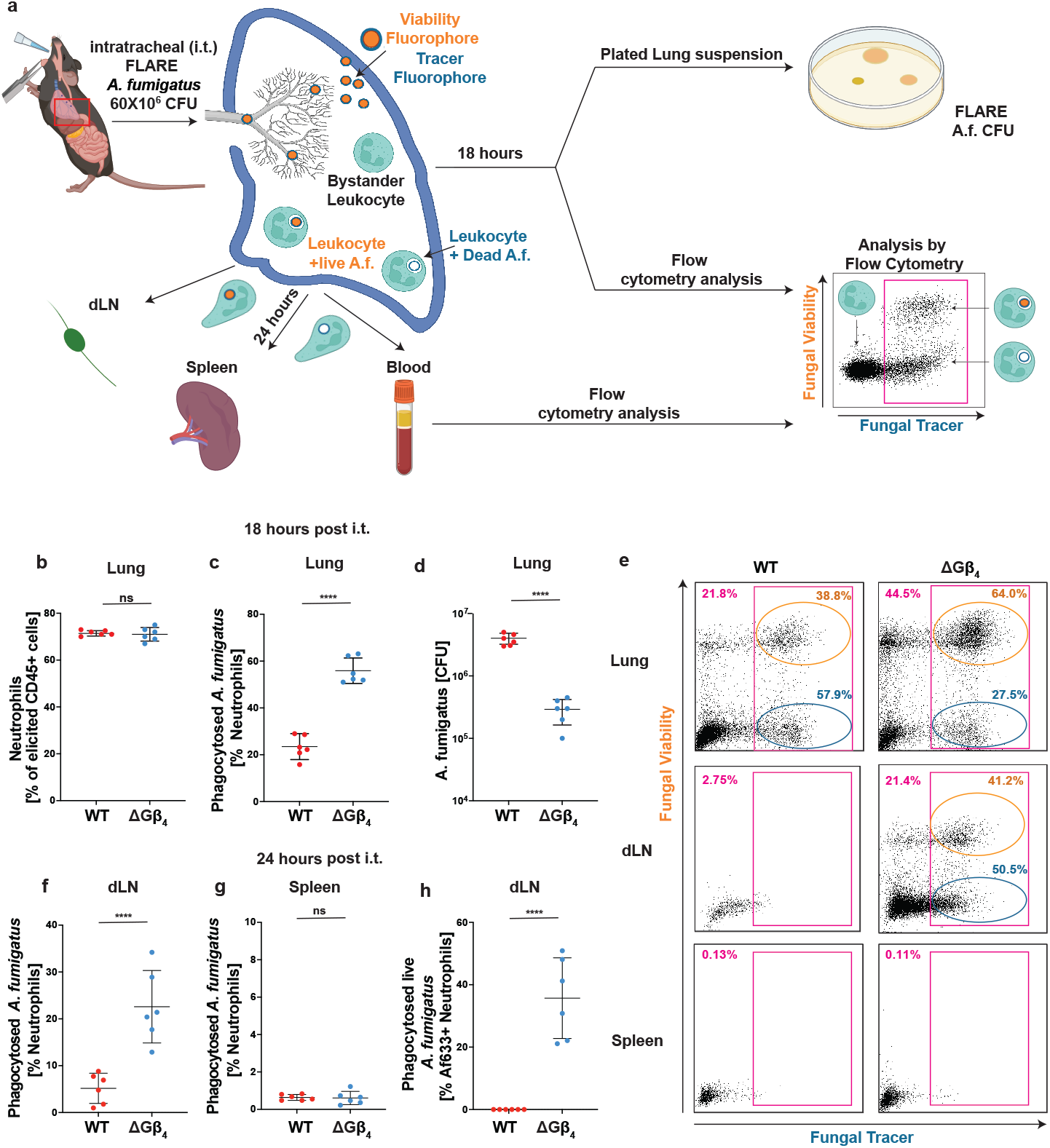
Gβ4 deficiency boosts phagocytic responses during in vivo fungal infection. WT and Gnb4-/- mice infected with FLARE A. fumigatus were assessed for in vivo phagocytosis and fungal infection after 18 h or 24 h. a, Schematic of the FLARE system. Conidia express dsRed and are labeled with AF633. Phagocytes become dsRed+AF633+ upon conidial uptake, and then transition to dsRed-AF633+ as internalized conidia are destroyed. b, Neutrophil infiltration into the lung, 18 h post-infection. c, Quantification of A. fumigatus phagocytosis by lung neutrophils at the 18 h time point. d, Fungal CFU in lung extracts at the 18 h time point. Data in b-d represent mean s.d. of 3 experiments. Unpaired t test **** p< 0.0001. e, Representative flow cytometry plots showing neutrophil phagocytosis of FLARE A. fumigatus in the lungs, dLN, and spleen 24 h post-infection (magenta gates). Gates for neutrophils containing live (orange) and dead (blue) conidia are shown. f, Quantification of neutrophils bearing engulfed FLARE A. fumigatus in the dLN, 24 h post-infection. g, Quantification of neutrophils bearing engulfed FLARE A. fumigatus in the spleen, 24 h post-infection. h, Quantification of the neutrophils in the dLN containing live A. fumigatus, 24 h post-infection. Data in f-h represent mean s.d. of 3 experiments. Unpaired t test **** p< 0.0001.

Given that Gβ4 deficient HL-60 cells failed to arrest their motility after cargo uptake (Fig. 2b-e), we speculated that Gnb4-/- phagocytes might exhibit altered trafficking behavior in vivo. Accordingly, we examined the organ distribution of phagocytic WT and Gnb4-/- neutrophils 24 hours after FLARE infection (Fig. 6a). In WT mice, neutrophils bearing AF633-labeled material were found predominantly in the lungs, with trace amounts in the draining lymph nodes (dLNs) and essentially none in the spleen and blood (Fig. 6e-g, fig. S9d). These observations were consistent with the expectation that phagocytosis inhibits motility after conidial uptake in the lungs, thereby preventing the dissemination of neutrophils with internalized pathogens to other organs. By contrast, we observed a substantial number of AF633+ neutrophils in the dLNs of Gnb4-/- animals (Fig. 6e-f), suggesting that Gβ4 deficiency at least partially relieves the brake on motility applied by phagocytosis. Importantly, whereas essentially all AF633+ neutrophils in WT dLNs were DsRed-, indicating that they had degraded their internalized conidia, a substantial fraction of DsRed+AF633+ neutrophils were observed in Gnb4-/- dLNs (Fig. 6h). Given that Gβ4 deficiency does not affect phagolysosome maturation (Fig. 1), these results strongly suggest that Gnb4-/- neutrophils continue to migrate after conidial uptake, enabling them to reach proximal organs like the dLN before their cargo is broken down. We conclude that Gβ4 deficiency not only increases the phagocytic capacity of myeloid cells in vivo but also disrupts the cross-regulatory relationship between phagocytosis and migration.

## Discussion

Taken together, our results identify Gβ4 as a critical regulator of plasma membrane abundance in myeloid cells and demonstrate that the plasma membrane exerts biophysical control over phagocytic capacity and functional crosstalk. Because plasma membrane must be mobilized to build pro-trusive cellular structures, limiting its abundance provides a simple mechanism for not only constraining the magnitude of a given architectural response (e.g. phagocytosis) but also enabling cross-regulation between responses (e.g. phagocytosis inhibiting migration). This scarcity model is consistent with prior work documenting antagonism between uptake behaviors, such as phagocytosis and micropinocytosis, and migration(Bretou et al. 2017; Chabaud et al. 2015; Luo et al. 2009). Thus, the mechanisms governing plasma membrane allocation in myeloid cells effectively dictate their functional potential. Notably, the plasma membrane dependent control mechanism studied here appears to affect most strongly the phagocytosis of large (10 um diameter) cargos, such as apoptotic cells and cancer cells, while having less impact on the uptake of small entities like bacteria. This distinction likely reflects the fact that engulfing large, unbroken targets with high surface area places a disproportionate bur-den on plasma membrane mobilization. The importance of Gβ4 for regulating this type of phagocytosis suggests that it may be particularly relevant for processes that involve the clearance of large eukaryotic cells, such as tissue homeostasis and anti-tumor immunosurveillance. The dramatic functional gains exhibited by Gb4 neutrophils and macrophages raise the question of why Gβ4 dependent control of membrane abundance evolved in the first place. One obvious answer is that tight cross-inhibition between phagocytosis and migration is essential for immune function, at least in some contexts. Indeed, we found that Gnb4-/- neutrophils not only take up more A. fumigatus conidia in vivo but also traffic them out of the lung to the dLN. While increased fungal dispersion did not appear to compromise immunity in these experiments, one could imagine that a pathogen more capable of resisting or escaping the phagolysosome could exploit the dysregulated migration of Gβ4 deficient phagocytes to spread. Although we cannot at present rule out a role for Gβ4 in the acute regulation of phagocytosis and motility, we favor a model in which the Gβ4 pathway acts during myeloid cell differentiation to dictate the size and composition of the plasma membrane, which then serves as a master mechanoregulator of morphology and effector responses in the terminally differentiated state (fig. S10). This model is consistent with our observations that Gβ4 neutrophils and macrophages exhibit architectural abnormalities (Fig. 3 and fig. S4) and also that acute myriocin treatment fails to reverse the Gb4 phagocytosis phenotype (fig. S5f). We postulate that differential Gb4 expression among myeloid progenitors might promote functional diversification by enabling the formation of differentiated subsets with a spectrum of plasma membrane-defined phagocytic and migratory setpoints. This type of diversity could enable the innate immune system to engage effectively with a wide variety of homeostatic and microbial challenges. Gβ4 mutations have been associated with Charcot-Marie-Tooth disease (CMTD)(Lassuthova et al. 2017; Soong et al. 2013; Hsu et al. 2019; Kwon et al. 2021), a hereditary neurological disorder characterized by the progressive demyelination of peripheral nerves. Although the pathogenesis of this disease is generally thought to arise from cell-intrinsic glial dysfunction, macrophages routinely patrol peripheral nerves and are well-positioned to induce autoimmune neuropathy(Zigmond and Echevarria 2019; Kolter, Kierdorf, and Henneke 2020). Interestingly, macrophage depletion has been shown to attenuate neurodegeneration in mouse CMTD models(Klein, Patzko, et al. 2015; Klein, Groh, et al. 2015; Carenini et al. 2001). In light of these reports, our data raise the intriguing possibility that dysregulated macrophage phagocytosis might contribute to at least some forms of CMTD. The identification of Gb4 signaling and sphingolipid synthesis as key regulators of phagocytic capacity in myeloid cells reveals heretofore unexplored avenues for enhancing innate immunity in therapeutic contexts. By targeting the architectural basis of cargo uptake, one could potentially modulate phagocyte activity in a manner that is agnostic to specific targets. We anticipate that an approach like this would be particularly useful for treating systemic microbial infections and for enhancing the anti-tumor potential of chimeric antigen receptor (CAR)-macrophages(Sloas, Gill, and Klichinsky 2021; Alvey and Discher 2017). Exploring these possibilities in additional translational experimental systems will be an interesting topic for future research.

## ACKNOWLEDGEMENTS

The authors thank the Molecular Cytology Core Facility, the Cell Metabolism Core Facility, the Flow Cytometry Core Facility, and the Integrated Genomics Operation at MSKCC for technical support; the New York Structural Biology Center for assistance with FIB-SEM; and members of the M.H., O.D.W., and J.G.C. labs for advice. Supported in part by the NIH (R01-AI087644 to M.H., R35-GM118167 to O.D.W., P30-CA008748 to MSKCC), the National Science Foundation (2019598 and 1548297 to O.D.W.), the Schmidt Science Fellows Program (B. Y. W.), the Cancer Research Institute (B. Y. W.), and the MSKCC Basic Research Postdoc Innovation Award (B. Y. W).

## AUTHOR CONTRIBUTIONS

B.Y.W, J.G.C., O.D.W., and M.H. conceived of the project, analyzed data, and wrote the manuscript. B.Y.W, A.H.S., A.Y., C.J., T.L, M.T., M.S., A.S., A.L.S., M.G., J.C., T.Z. and H.A.P. performed experiments or analysis for the manuscript. Y.Y.W made schematic figures for the manuscript. F.G, L.C.K., J.S.A.P, and T.H. provided reagents and helpful expertise.

## Supplement

**Supplementary Movie 1: Gβ4 deficiency enhances the phagocytic activity of HL-60 cells. (related to Fig. 1E)** Gβ4 deficiency enhances the phagocytic activity of HL-60 cells. WT and ΔGβ4 HL-60 cells expressing F-tractin (red) were mixed with IgG coated MPs (green) and imaged by spinning disk confocal microscopy. Video shows representative interactions between MPs and the indicated HL-60 cells, viewed at 15× time-lapse. ΔGβ4 HL-60 cells engulf targets much more quickly than do WT controls.

**Supplementary Movie 2: Gβ4 attenuates cell migration and alters crosstalk between phagocytosis and motility. (related to Fig. 2)** ΔGβ4 and WT cells were seeded onto a fibronectin coated slide and exposed to 200 nM fMLF delivered through a micropipette over the course of an hour. The first video compares the motility of ΔGβ4 and WT cells prior to phagocytosis, viewed at 280× time-lapse. The second video shows a mixture of ΔGβ4 and WT cells migrating after uptake of one IgG-coated MP, viewed at 280× time-lapse. ΔGβ4 cells contain green MPs, while WT cells contain red MPs. AF647 was added to the fMLF in all experiments to facilitate visualization of the fMLF gradient. Time in MM:SS is indicated in the upper left corner of each video.

**Supplementary Movie 3: FIB-SEM analysis of** D**Gβ4 and WT HL-60 cells. (related to Fig. 3b-e)** ΔGβ4 and WT HL-60 cells were stained with potassium ferrocyanide and osmium tetroxide and imaged by FIB-SEM. 3-D reconstructions of representative ΔGβ4 and WT cells are shown, with plasma membrane shown in semi-transparent green and lipid droplets in solid magenta.

**Fig. S1.**
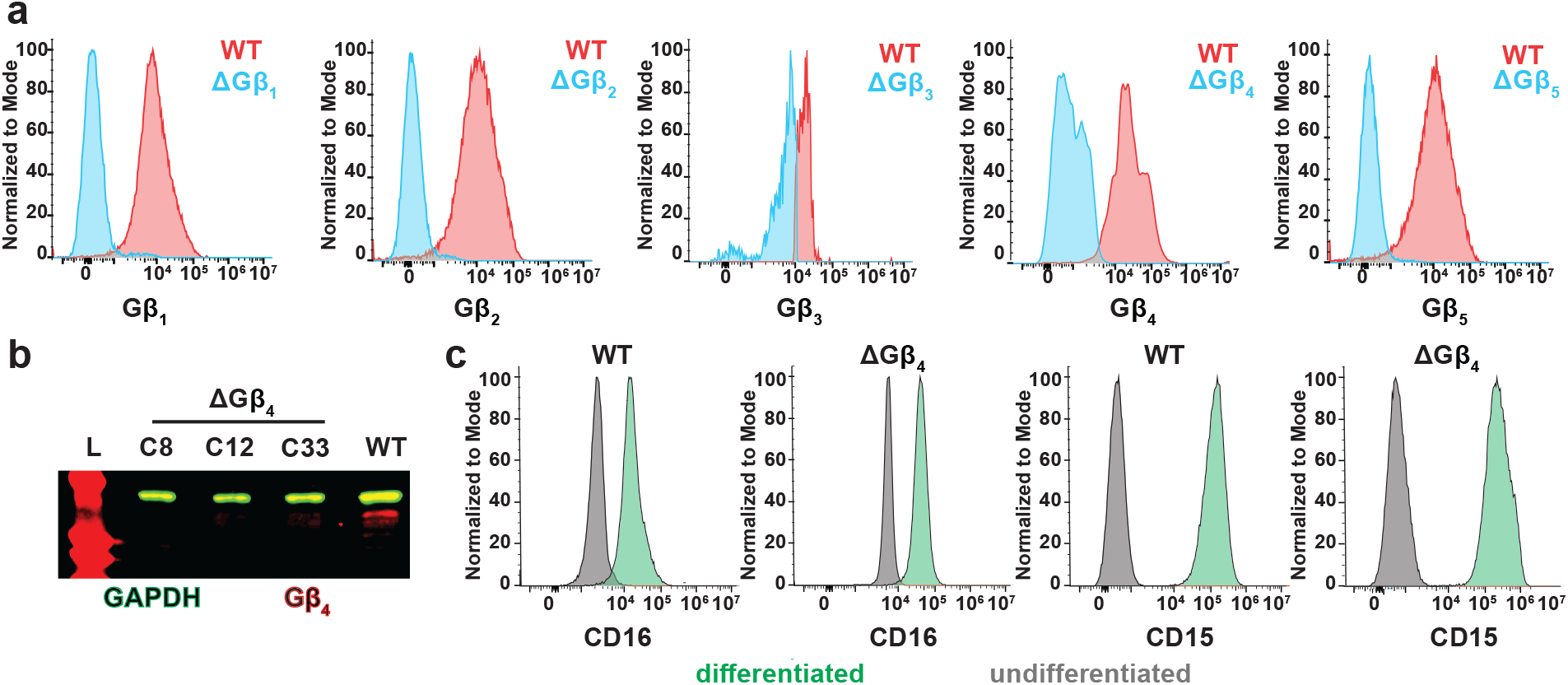
Deletion of Gβ 4 does not affect differentiation of HL-60 progenitors into neutrophil-like cells. a, Flow cytometric validation of the indicated Gβ knockouts. Histograms show WT cells in red and each Gβ knock-out cell line in blue. b, Immunoblot analysis of Gβ 4 (red) in WT and ΔGβ 4 HL-60 cells. GAPDH (green) served as a loading control. c, WT and ΔGβ 4 HL-60 progenitor cells were differentiated into neutrophils and stained for CD15 or CD16. In each histogram, differentiated cells are shown in green and progenitors in gray.

**Fig. S2.**
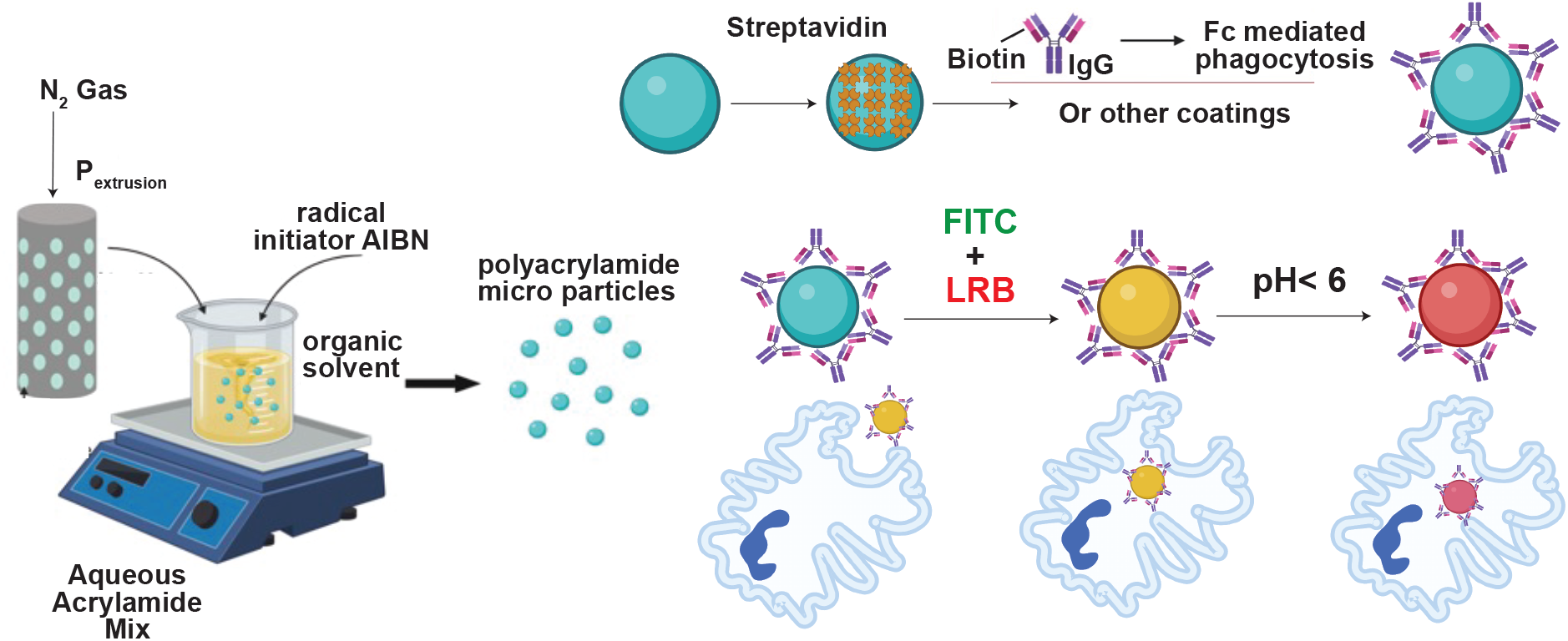
Preparation of MP cargo. Schematic diagram showing fabrication of polyacrylamide MPs, conjugation with phagocytic ligands (e.g. IgG), and labeling with LRB and FITC. After MP uptake, acidification of the phagolysosome quenches FITC fluorescence, turning MPs from yellow into red.

**Fig. S3.**
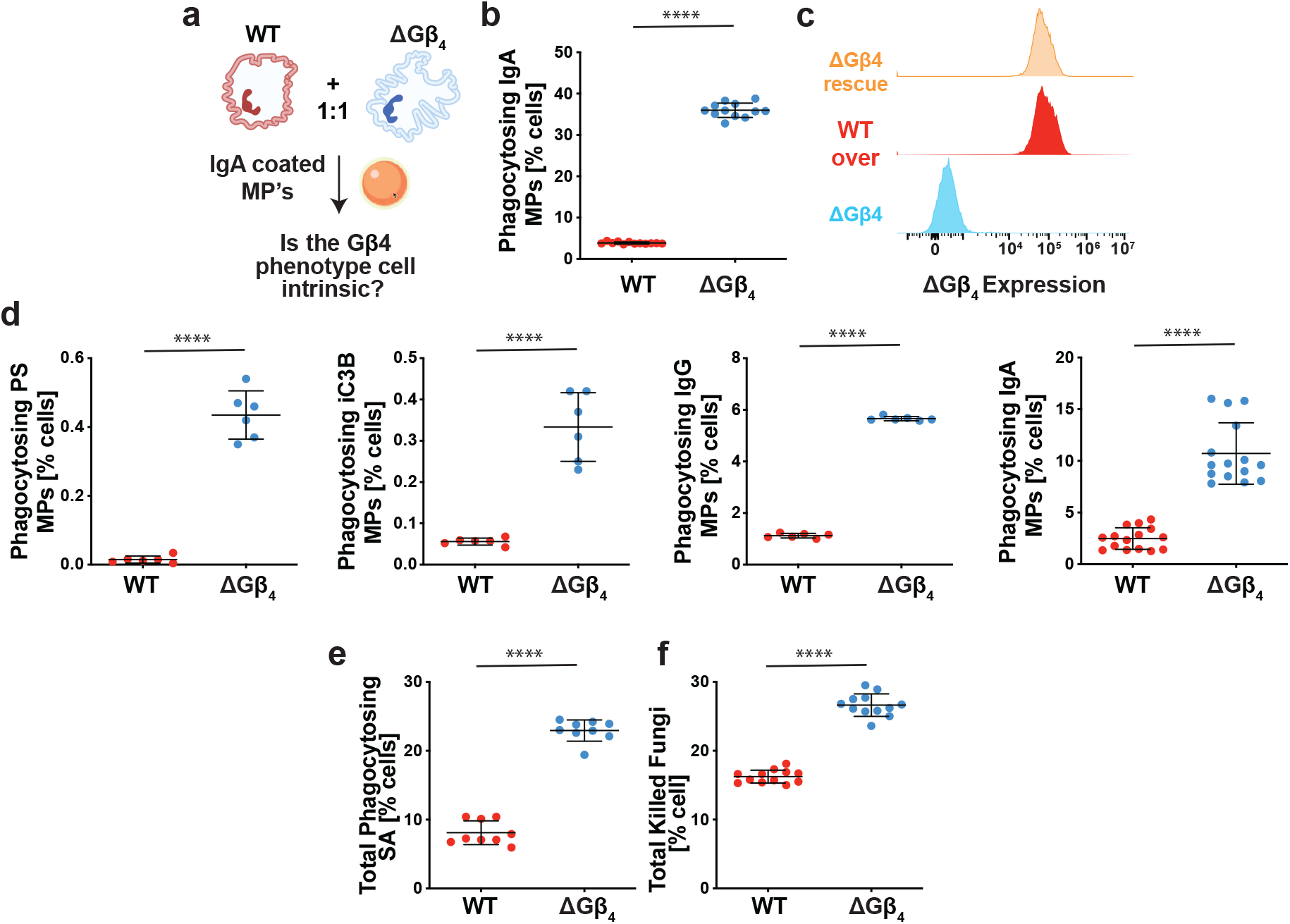
ΔGβ4 hyperphagy applies to a wide range of phagocytic targets. a-b, ΔGβ4 and WT HL-60 cells were stained with different fluorophores, mixed 1:1, and then challenged with IgG coated MPs. a, Schematic of the experimental design. b, Quantification of phagocytic uptake after 3 h. Data are mean s.d. of three experiments. Unpaired t test **** p< 0.0001. c, WT and ΔGβ4 HL-60 cells were lentivirally transduced with exogenous Gβ4 (WT over and ΔGβ4 rescue). Histograms show Gβ4 expression in each cell line, relative to ΔGβ4 HL-60 cells. d, ΔGβ4 and WT HL-60 cells were challenged with MPs coated with phosphatidylserine (PS), complement (iC3B), human IgG, or human IgA to mimic efferocytosis, complement, Fc γ R, or Fc α R mediated phagocytosis, respectively. Phagocytosis was quantified after 3 h. Data are mean s.d. of 3 experiments. Unpaired t-test **** p< 0.0001. e, ΔGβ4 and WT HL-60 cells were challenged with S. aureus (SA) bioparticles, and phagocytosis was quantified after 1 h. f, ΔGβ4 and WT HL-60 cells were challenged with Aspergillus fumigatus fungal spores, and fungal killing (DsRed degradation) quantified after 1 h. Killing frequency was determined by dividing the number of HL-60 cells containing dead conidia (dsRed-AF633+) by the total number of AF633+ cells. In e and f, Data are mean s.d. of three experiments. Unpaired t test **** p< 0.0001.

**Fig. S4.**
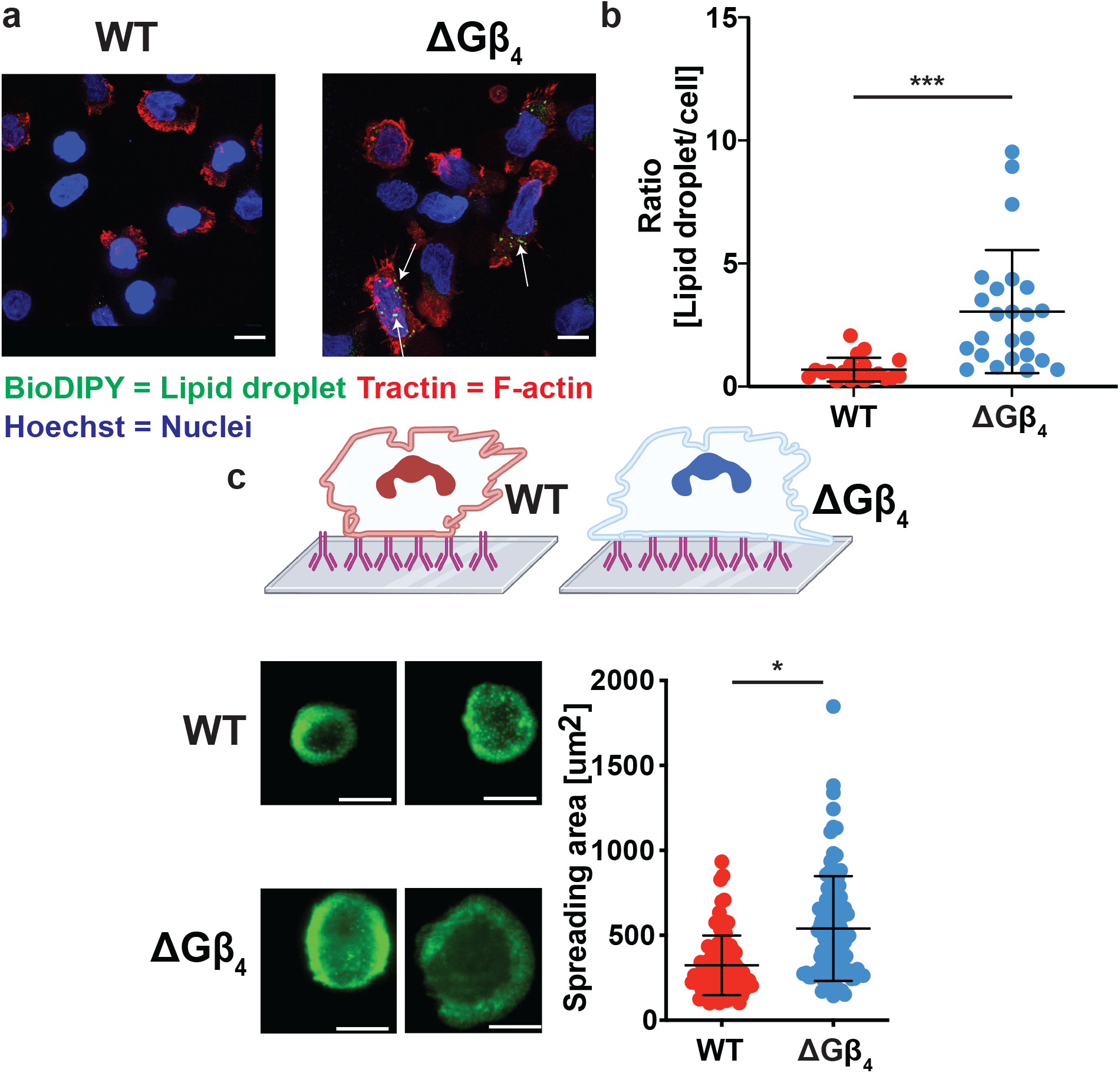
ΔGβ4 cells have excess plasma membrane and lipid droplets. a-b, ΔGβ4 and WT HL-60 cells expressing F-tractin-mCherry were stained with BioDIPY 493/503 and Hoechst to visualize lipid droplets and nuclei, respectively. a, Representative images, with white arrows indicating lipid droplets in ΔGβ4 cells. Scale bars = 7 μm. b. Quantification of lipid droplet content per cell, derived from 20 frames. Unpaired t test *** p< 0.001. c, ΔGβ4 and WT HL-60 cells expressing F-tractin-GFP were stained with cell mask orange and applied to IgG coated glass coverslips. Above, schematic diagram of the assay. Below left, TIRF images of representative cells after frustrated phagosome formation. Scale bars = 5 μm. Below right, quantification of phagocytic cup size. Data are mean s.d. of three experiments, WT (n=45 cells), ΔGβ4 (n= 36 cells). Unpaired t-test * p< 0.05.

**Fig. S5.**
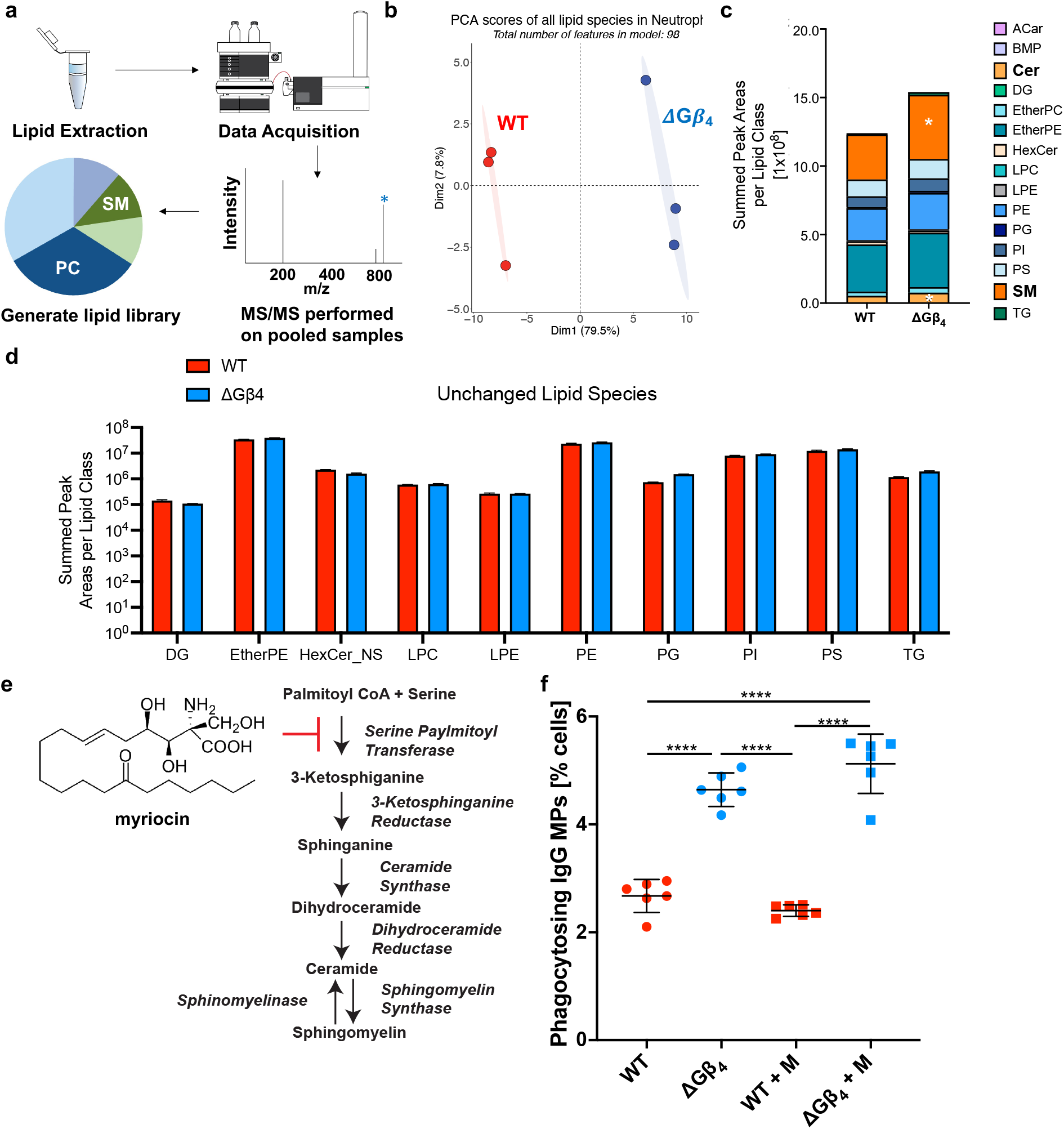
ΔGβ4 cells are enriched in sphingolipids. a-d, ΔGβ 4 and WT HL-60 cells were subjected to comparative lipidomics at steady state. a, Schematic of the lipidomic pipeline. b, PCA analysis of ΔGβ4 and WT biological replicates. A total of 98 different lipid species were identified. c, Cumulative bar graph showing the distribution of lipids in each experimental group. High abundance phosphatidylcholine (PC) species have been removed from the graph for clarity. Sphingomyelin (SM) and ceramide (Cer) contributions are indicated by white asterisks. d, Bar graph showing the lipid species that are unchanged between samples or over-represented in WT cells. e, Schematic of myriocin inhibition of sphingolipid synthesis. f, Differentiated ΔGβ4 and WT HL-60 cells were treated with myriocin (M) acutely for 12 h and then were challenged with IgG coated MPs. Phagocytosis was quantified 3 h after MP challenge. Data are mean s.d. of 3 experiments, WT (n=6), ΔGβ4 (n= 6). One-way ANOVA, **** p< 0.0001.

**Fig. S6.**
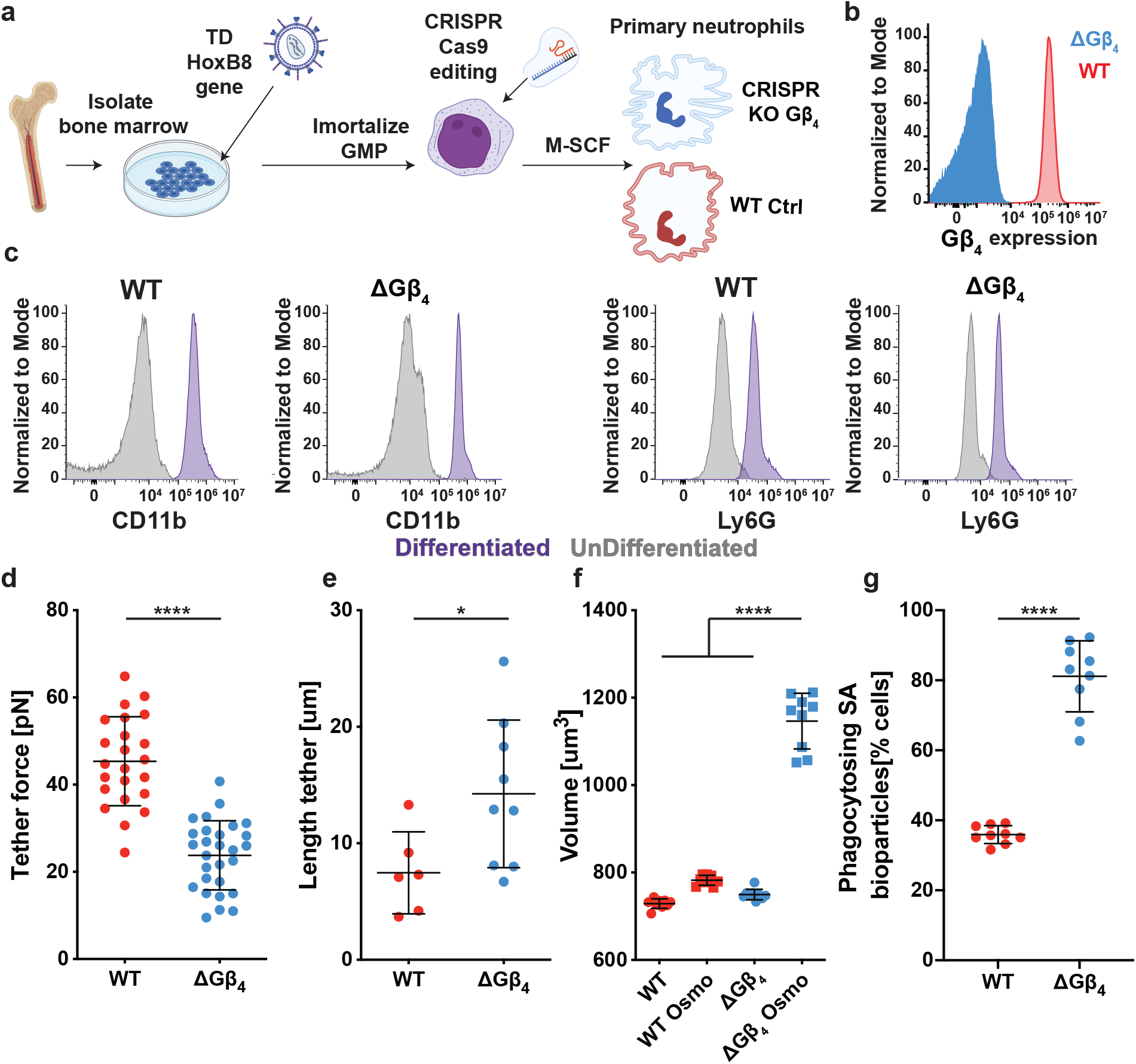
ΔGβ4 primary murine neutrophils derived from ER-HoxB8 GMPs exhibit excess plasma membrane and hyperphagy. a, Schematic showing the generation of ΔGβ4 and WT control primary murine neutrophils from ER-HoxB8 progenitors. b, Flow cytometric validation of Gβ4 deletion in ER-HoxB8 neutrophils. c, Flow cytometric analysis of CD11b and Ly6G expression in the indicated ER-HoxB8 progenitors (gray) and differentiated neutrophils (purple). d-e, Membrane tethers were generated from WT and ΔGβ4 ER-HoxB8 neutrophils using an optical trap. d, Quantification of membrane tension. e, Quantification of membrane tether length. Data in d and e are mean s.d. of 3 experiments. Unpaired t-test, * p<0.05, **** p< 0.0001. f, Volume of WT and ΔG4 ER-HoxB8 neutrophils determined before and after osmotic shock (osmo). Data are mean s.d. from 3 experiments. One-way ANOVA **** p< 0.0001. g, ΔGβ4 and WT ER-HoxB8 neutrophils were challenged with pHrodoRed stained S. aureus bioparticles, and phagocytosis was quantified after 1 h. Data are from 3 separate experiments. Unpaired t test **** p< 0.0001.

**Fig. S7.**
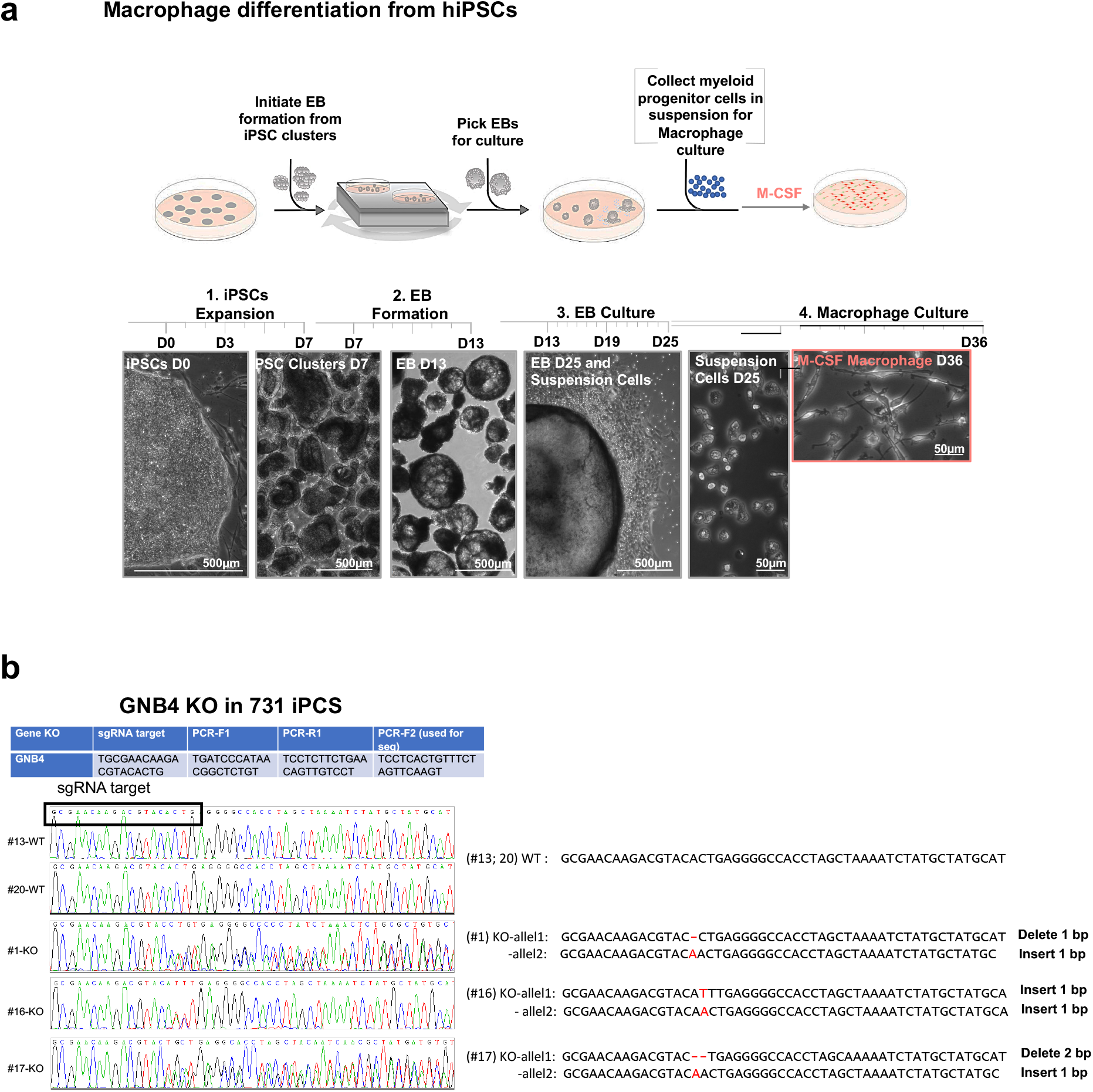
Generation of ΔGβ4 primary human macrophages from hiPSCs. a, ΔGβ4 and isogenic WT control hiPSCs were first plated onto irradiated mouse embryonic fibroblasts. This led to the formation of embryoid bodies (EBs), which continuously produced granulocyte monocyte progenitor (GMPs). GMPs were isolated and differentiated into primary human macrophages over the course of 3 to 5 days. b, Above, sgRNAs used for CRISPR-Cas9 targeting, along with the oligos used to amplify and sequence the targeted GNB4 locus. Below left, Sanger sequencing plots of the targeted region from 2 WT and 3 ΔGβ4 (KO) clones. Below right, interpreted sequences from each sequencing reaction. Note the presence of indels on both GNB4 alleles in all three KO samples.

**Fig. S8.**
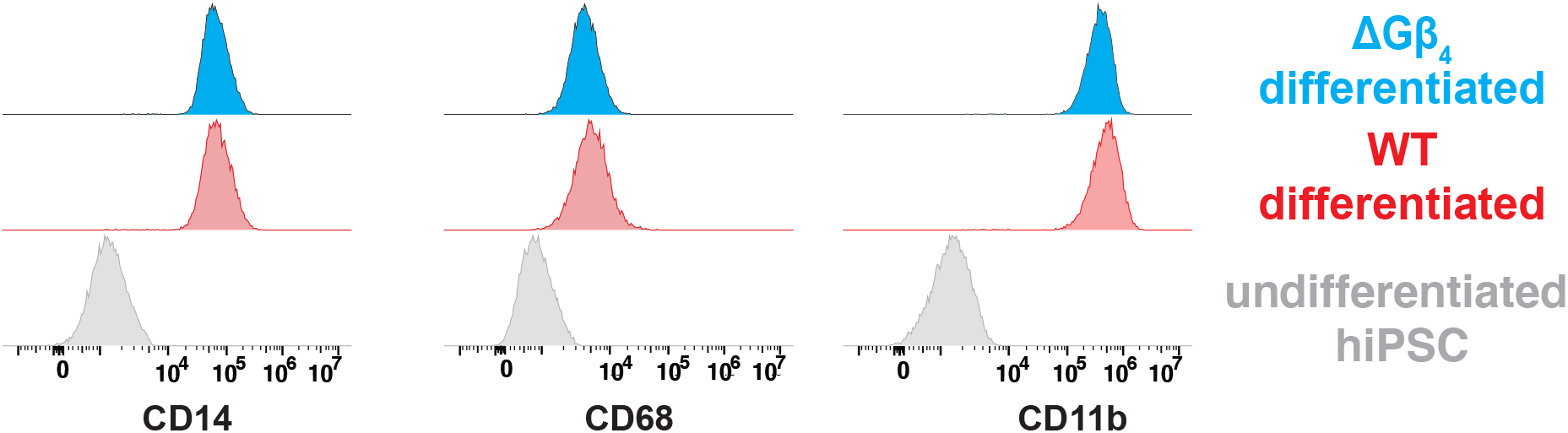
Gβ4 deficiency does not block macrophage differentiation from hiPSCs. a, CD14, CD68, and CD11b expression on undifferentiated hiSPCs (gray) and on ΔGβ4 (blue) and WT control (red) hiPSC-derived macrophages.

**Fig. S9.**
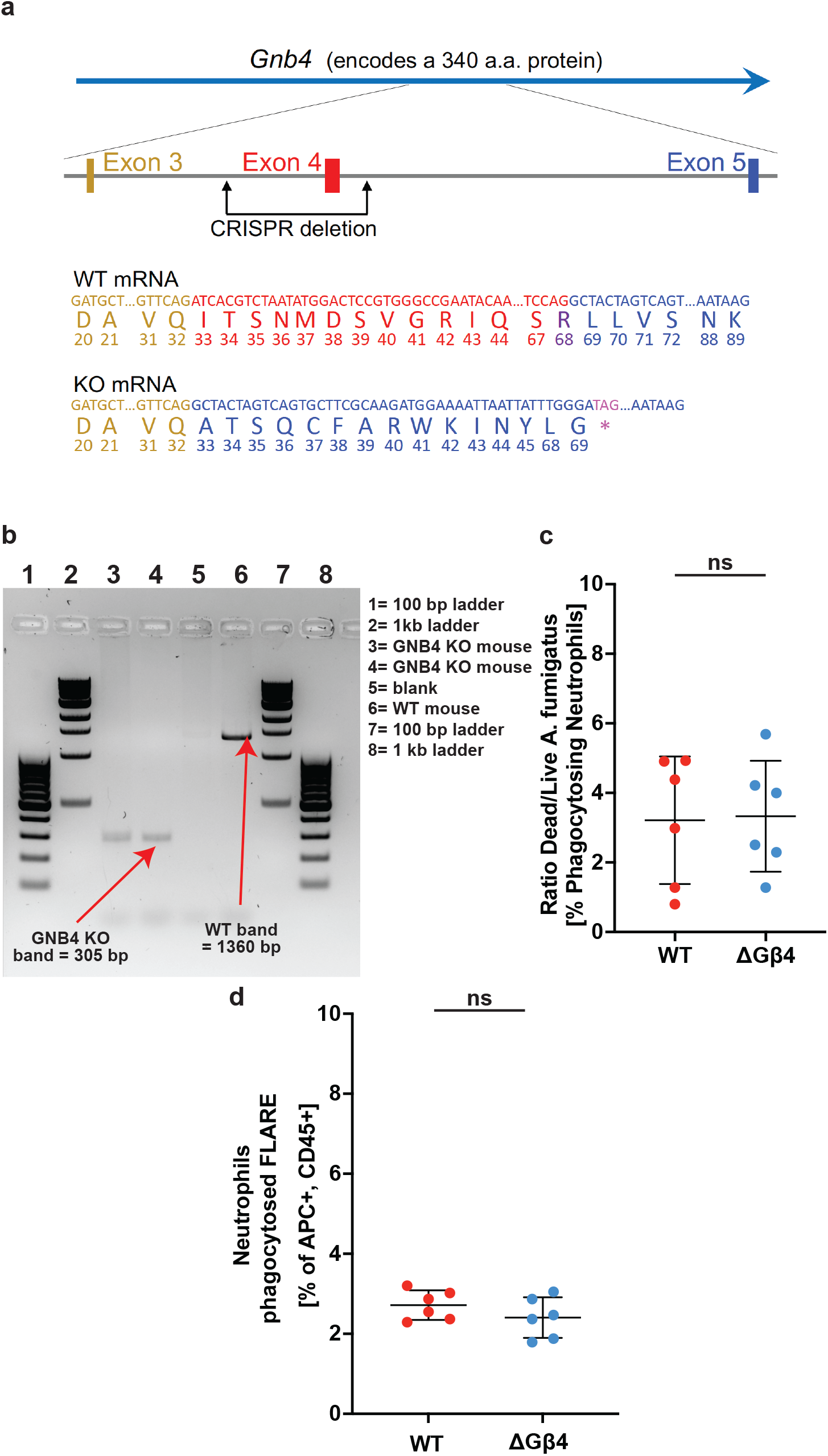
Generation of Gnb4-/- mice. a-b, Gnb4-/- mice were created by targeting exon 4 using CRISPR-Cas9. a, Above, a schematic diagram indicating the two double stranded breaks induced by CRISPR-Cas9, which afforded deletion of exon 4 along with some associated intronic sequences. Below, predicted mRNA and protein generated by the WT and knock-out Gnb4 loci. Only relevant sequences from exons 3, 4, and 5 are shown. Deletion of exon 4 leads to translational frameshift and premature termination of the Gnb4 protein at the magenta stop codon. b, Representative genotyping showing Gnb4 knock-out (KO) and WT bands. c, WT and Gnb4-/- mice infected with FLARE A. fumigatus were assessed for in vivo phagocytosis and fungal infection after 18 h. Graph shows quantification of fungal killing frequency, determined by dividing the number of Ly6G+CD11b+Siglec F-neutrophils containing dead conidia (dsRed-AF633+) by the total number of AF633+ neutrophils. Data are mean s.d. from 3 experiments. d, Quantification of AF633+ neutrophils in the blood 24 hours post i.t. infection with FLARE conidia.

**Fig. S10.**
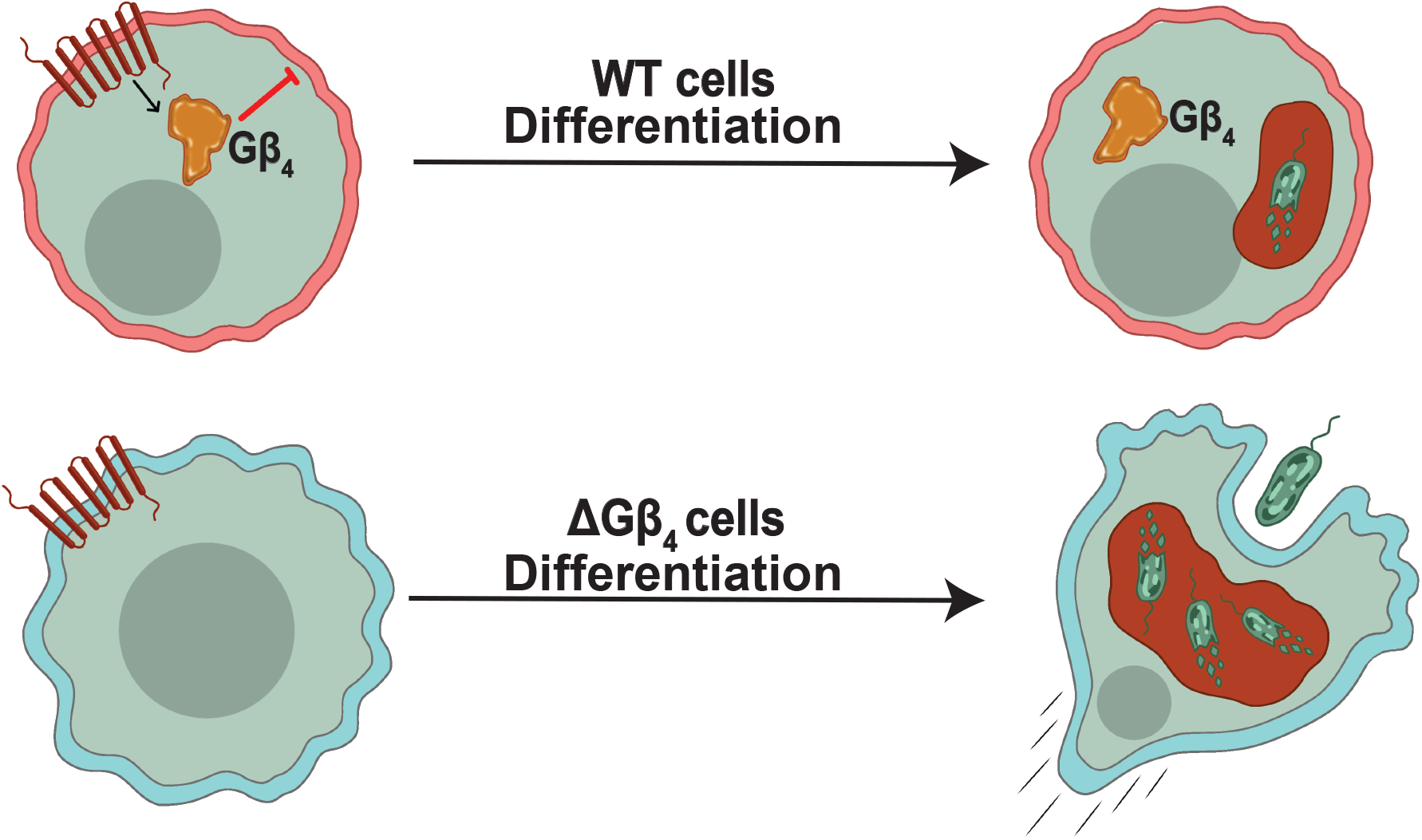
Model for Gβ4-mediated control of myeloid effector function. Gβ4 signaling acts to restrain plasma membrane production during the differentiation of neutrophils and macrophages. This curtails phagocytic potential but enables effective cross-regulation between phagocytosis and migration. In the absence of Gβ4, myeloid cells acquire excess plasma membrane, enabling them to engulf substantially more cargo while continuing to migrate in search of additional targets.

## Experimental details

### Cell lines, growth, and differentiation conditions

HL-60 cells were obtained from the laboratory of Henry Bourne and validated by RNA sequencing. Jurkat human T cell lymphoblasts (clone E6-1) were purchased from ATCC (cat TIB-152). RAMOS B cell Burkitt’s Lymphoma cells were purchased from ATTC (cat CRL-1596). HL-60, Jurkat, and RAMOS cells were grown in RPMI media supplemented with L-glutamine, 25 mM Hepes, 10 percent (vol/vol) heat inactivated fetal bovine serum (FBS), at 1 percent Pen/Strep. HL-60 progenitor cells were split to 2 × 105 cells/mL every 2 days and grown at 37 ºC, 5 percent CO2. HL-60 progenitors were differentiated into neutrophil-like cells by transfer into 1.5 percent (vol/vol) DMSO at 2 × 105 cells/mL followed by 5 days of incubation at 37 ºC, 5 percent CO2. HEK293T lentiX cells (Takara Bio, cat 632180) were grown in DMEM containing 10 percent (vol/vol) heat-inactivated FBS and 1 percent Pen/Step and maintained at 37 ºC, 5 percent CO2. All cell lines were routinely checked for mycoplasma contamination.

### HL-60 manipulation

Gβ 1-5 were deleted from HL-60 progenitors using the IDT Alt-R CRISPR-Cas9 system (IDT). Briefly, crRNAs specific for Gβ 1-5 (see table 1 below for sequence details) were resuspended in sterile Duplex Buffer (IDT) to a final concentration of 200 μM. Each crRNA was then mixed with tracrRNA-ATTO 550 (IDT) at a 1:1 ratio and annealed using a thermocycler (BioRad). sgRNA:Cas9 RNP’s were formed by mixing 0.3 μL of 36 μM Cas9 protein (IDT) with annealed sgRNAs at a 1:1 ratio followed by room temperature incubation for 10-20 minutes. HL-60 progenitor cells at a density of 2.2 × 106 (>95

Table 1: sgRNA RNP sequences crRNA specific for Gβ s sequence Hs.Cas9.GNB1.1.AA TGAGCTTGACCAGTTACGGC Hs.Cas9.GNB2.1.AF TCTTTGCCAGGTGCCCACGG Hs.Cas9.GNB3.1.AD TGCCAGAGTAACGTCAGCAC Hs.Cas9.GNB4.1.AA GC-GAACAAGACGTACACTGA Hs.Cas9.GNB5.1.AG CGAGAGCCTCAAGGGCAAGC Hs.Cas9.GNB4.1.AB ATAAGCACAGGTCAT-CACCC

Table 2: Rabbit monoclonal antibodies used for staining G-Beta Supplier Ref GNB1 Abcam ab137635 GNB2 Abcam ab108504 GNB3 Abcam ab154866 GNB4 Abcam ab223113 GNB5 Abcam ab185206

To overexpress Gβ4, a GNB4 gene block (IDT) containing homology arms to the lentiviral vector pLVX-Puro (CloneTech) was inserted into XhoI-digested pLVX-Puro using the In-Fusion HD Kit (Takara Bio). The homology arms were 5’-ctaccggactcagatctcga-3’ at the N-terminus of GNB4 and 5’-tcgagctcatcgggatcccgctcgacta-3’ at the C-terminus. The resulting plasmid was combined with plasmids encoding

(1) a minimal HIV pLVX with GNB4-2A-Puro transgene, (2) gag-pol from HIV53 and (3) appropriate viral glycoproteins (VSV-G), and the mixture then transfected into HEK293T lentiX cells (Takara Bio) using the Xtremegene transfection reagent (Roche Applied Science). Lentivirus was collected 24 and 48 hours post transfection and stored at -80 ºC until use. For transduction, 0.5 to 1 × 106 HL-60 progenitor cells were mixed with lentivirus preparations (1 mL of virus supplemented with polybrene and HEPES) and centrifuged at 1400 × g at 37 ºC for 2 hours in either 24-well or 6-well polystyrene plates (Corning). Subsequently, 1 mL of RPMI complete media (10 percent FBS, 1 percent Pen/Strep) was added to the cells, followed by overnight incubation at 37 ºC, 5 percent CO2. Cells were then placed in 0.5 μg/mL puromycin for 14 days to select for successfully transduced cells. HL-60 progenitors transduced with lentivirus derived from empty pLVX-Puro were prepared in parallel as controls. To express F-tractin-mCherry in HL-60 cells, DNA encoding the F-tractin-mCherry fusion was amplified from a C1-F-tractin-mCherry plasmid (Addgene, 155218) and then subcloned into the lentiviral vector pLVX-M-Puro (CloneTech) by Gibson reaction. Lentivirus production and transduction was performed as described above.

### Fungal/bacterial lines

FLARE Aspergillus fumigatus conidia were prepared as previously described(Jhingran et al. 2012). Briefly, Af293 conidia were transformed with pAN7-1(Punt et al. 1987) and pPgpd-dsRed(Mikkelsen et al. 2003), and 5 × 108 of the resulting Af293-dsRed conidia were incubated with 0.5 mg/mL Biotin XX, SSE (B-6352; Invitrogen) in 1 mL of 50 mM carbonate buffer (pH 8.3) for 2 h at 4 ºC, washed once with 0.1 M Tris-HCl (pH 8), and then labeled using 0.02 mg/mL AF633-streptavidin (S-21375; Invitrogen) in 1 mL PBS for 30 min at room temperature. Double-labeled FLARE conidia were resuspended in PBS + 0.025 percent Tween 20 for experimental use. Pseudomonas aeruginosa expressing GFP (P. aeruginosa PA14 PrhlAB::gfp, kind gift from Joao Xavier) were labeled with AF633 by first incubating 5 × 107 bacterial cells with 0.5 mg/mL Biotin XX, SSE (B-6352; Invitrogen) in 1 ml 50 mM carbonate buffer (pH 8.3) for 2 h at 4 ºC. After washing with 0.1 M Tris-HCl (pH 8), cells were labeled using 0.02 mg/mL AF633-streptavidin (S-21375; Invitrogen) in 1 mL PBS for 30 min at RT, followed by resuspension in PBS + 0.025 percent Tween 20.

### Polyacrylamide microparticle (MP) cargo

10 μm diameter polyacrylamide MPs were synthesized as previously described(Vorselen et al. 2020), with modifications. Aqueous acrylamide solutions were prepared containing 150 mM MOPS (pH 7.4), 0.3 percent (v/v) tetramethylethylenediamene (TEMED), 150 mM NaOH, and 10 percent (v/v) acrylic acid. The mass fraction of total acrylamide was kept constant at 10 percent and the cross-linker mass fraction was 2.4 percent. This mixture was sparged with nitrogen gas for 15 minutes and then kept under nitrogen pressure. Hydrophobic Shirasu porous glass (SPG) filters (1.9 μm pore size) were sonicated under vacuum in n-dodecane, mounted on an internal pressure micro kit extruder (SPG Technology Co), and submerged into the organic phase, consisting of 99 percent hexanes and 1 percent (v/v) Span-80 (Sigma-Aldrich, S6760), with continuous stirring at 300 rpm. 10 mL of the acrylamide mixture was then loaded into the apparatus and extruded under nitrogen pressure into 250 mL of the hexane mixture. To initiate polymerization, the resulting emulsion was heated to 60 ºC and 2,2’-azobisisobutyronitrile (2 mg/mL, 500 mg total) was added. The reaction was kept at 60 ºC for 3 hours and then at 40 ºC overnight. Polymerized MPs were harvested by washing 3 times in hexanes and 2 times in ethanol (30 minutes, 4000 rpm), followed by resuspension in PBS. For functionalization, MPs were washed twice in activation buffer (100 mM MES, 20 mM NaCl, pH 6.0) and then incubated for 15 minutes with 40 mg/mL 1-ethyl-3-(3-dimethylaminopropyl) carbodiimide (EDC), 20 mg/mL N-hydroxysuccinimide (NHS) in activation buffer with 0.1 percent (v/v) Tween 20, vortexing briefly every 5 min. After 3 washes in PBST8 (PBS with 0.1 percent (v/v) Tween 20, adjusted to pH 8), MPs were resuspended in PBST8 and a protein/dye solution was added. For streptavidin labeling, 250 pmol streptavidin per million MPs was added. For acidification assays, MPs were conjugated with either 5-FITC-ethylenediamine (5 μM final, AAT Bioquest 126) and LissamineTM Rhodamine B Ethylene Diamene (5 μM final, Invitrogen L2424). After 2 h, unreacted NHS groups were blocked with ethanolamine (100 mM final, in 100 mM Tris HCl). Finally, MPs were washed 3 times in PBST, pH 7.4, and 3 times in PBS, pH 7.4. In certain phagocytosis experiments, labeled MPs were conjugated with pHrodo Dyes rather than FITC/LRB by first washing 3 times in 0.1M NaHCO3, pH 8.4, followed by a 1 h incubation in pHrodoTM Green STP Ester (Invitrogen P35369) or pHrodoTM iFL Red STP Ester (Invitrogen P36010) at 60 μM final concentration, followed by 3 washes in PBS, pH 7.4. Streptavidin conjugated MPs were loaded with Biotin-SP ChromPure Human IgG (Jackson ImmunoResearch 015-060-003), Biotin-SP ChromPure Human Serum IgA (Jackson ImmunoResearch 009-060-011), iC3B (Quidel A423), and Biotin-Phosphatidylserine (Echelon L-31B16) at 200 pmol per million MPs, 100 pmol per million MPs, 100 pmol per million MPs, and 50 pmol per million MPs, respectively, in 750 μL of PBS. iC3B was biotinylated prior to MP coating using the EZ-Link™ Sulfo-NHS-LC-Biotinylation Kit (ThermoFisher).

### Flow cytometric MP phagocytosis assay

Neutrophils stained with either Hoechst or CellVue Maroon (both from ThermoFisher) were seeded into fibronectin-coated 24 well plates at a density of 150,000 cells per well. FITC/LRB labeled MPs coated with a phagocytic target ligand (e.g. IgG) were then added at a 1:2 (MP:cell) ratio, followed by incubation at 37 ºC, 5 percent CO2 for 3 hours. Cells were then removed by trypsinization and transferred into FACS tubes for analysis using a Cytoflex LX machine (Beckman). Analysis was performed by first gating out free floating MPs and then identifying LRB+FITClow cells, indicative of successful uptake and acidification of MPs. In each sample, the total number of LRB+FITClow cells was normalized to the total number of live cells to generate a percent phagocytosis metric. In experiments using pHrodoGreen or pHrodoRed dyed MPs, the percent phagocytosis metric was calculated using pHrodoBright cells (in FITC or PE channels) as the numerator and total cells as the denominator. In experiments comparing Gβ4 and wild type HL-60 cells in the same sample, each cell type was labeled with a different dye (either Hoechst or CellVue Maroon, both from ThermoFisher) to distinguish them during endpoint flow cytometric analysis. Dyes were switched between experiments to control for potential effects of the dyes on phagocytosis.

### Widefield imaging of MP phagocytosis

Neutrophils or macrophages were stained using Hoechst and/or CellVue Maroon (both from ThermoFisher) and seeded into fibronectin coated 8-well chamber slides (ibidi) at a density of 100,000 or 50,000 cells per well, respectively. FITC/LRB labeled IgG coated MPs were then added at a 1:2 (MP:cell) ratio, and the samples imaged on a Zeiss Axiovert microscope using a 20× objective lens for at least 2 h at 1 min intervals. DAPI, FITC, and TRITC images were collected at each time point. Phagocytosed MPs were identified in Fiji by a reduction in FITC signal due to phagolysosome acidification. Percent phagocytosis was determined by dividing the number of phagocytes containing acidified MPs by the total number of phagocytes in each frame.

### High resolution imaging of phagocytic cup closure

ΔGβ4 and WT HL-60 cells expressing F-tractin-mCherry were stained with CellVue Maroon (ThermoFisher) using the manufacturer’s guidelines and then seeded at 100,000 cells per well into fibronectin coated 8-well ibidi chambers. IgG-coated, FITC-labeled MPs were either added at a 1:2 (MP:cell) ratio after cell seeding or, in some experiments, were attached to the bottom of the slide by first coating with biotin poly-L-lysine (ThermoFisher) and then adding MPs for attachment. Cells were imaged using a SoRa spinning disk confocal microscope (Nikon), a Stellaris spinning disk confocal microscope (Leica), or an LSM880 confocal microscope (Zeiss) fitted with 60× objective lenses. In general, time-lapse experiments encompassed 30 minutes total with FITC, TRITC, and Cy5 images collected every 15 seconds. Images were analyzed using Fiji and/or Imaris to assess phagocytic cup progression and completion. Cup initiation was determined by an enrichment of F-tractin adjacent to the MP. Phagocytic cup closure was defined as total MP encirclement by F-tractin/plasma membrane. Time to MP engulfment was determined by subtracting the time of cup closure from the time of initiation. Phagocytic cups that were not completed by the end of the 30 min time-lapse were considered “aborted”.

### Staphylococcus aureus bioparticle phagocytosis assay

HL-60 cells or primary neutrophils were seeded in fibronectin coated 24-well plates at 250,000 cells per well. 10 μL of S. aureus pHrodoRed bioparticles (ThermoFisher), resuspended in sterile PBS at a final concentration of 2 mg/mL, were then added to each well, followed by incubation at 37 ºC, 5 percent CO2, for 1 hour. Cells were then harvested and transferred into FACS tubes for analysis using a Cytoflex LX machine (Beckman). Percent phagocytosis was determined by dividing the total number of pHrodoRed+ cells by the total number of cells in each sample.

### Jurkat efferocytosis assay

5 × 106 Jurkat cells in 10 mL complete RPMI (10 percent FBS, 1 percent Pen/Strep) were transferred into a lidless 10 cm dish and subjected to 1500 microJ UV using a UV stratalinker (Marshall Scientific), followed by a 4 h incubation at 37 ºC, 5 percent CO2. The cell corpses were then resuspended in 10 mL of room temperature Hank’s buffered salt solution (HBSS), stained with either CypHer5E NHS Ester (1:4000 in 4 mL PBS, Cytiva) or pHrodoRed (1:5000, Invitrogen) for 10 min at 37 ºC, and then washed into complete RPMI (10 percent FBS, 1 percent Pen/Strep). Terminally differentiated HL-60 cells were stained with CFSE (1:1000, ThermoFisher) and then mixed with Jurkat corpses at a ratio of (1:2 Jurkat cell corpse/HL-60 cell) in a final volume of 1 mL, followed by incubation for 3 h at 37 ºC, 5Percent CO2. The cells were then harvested and transferred into FACS tubes for analysis using a Cytoflex LX machine (Beckman). Percent phagocytosis was determined by dividing the total number of FITC+Cypher+ or FITC+pHrodoRed+ HL-60 cells by the total number of HL-60 cells (FITC+) in each sample.

### Aspergillus fumigatus FLARE fungal phagocytosis and killing assay

HL-60 cells were seeded at 150,000 cells per well in a 24 well plate and then mixed with FLARE A. fumigatus (see procedure above for preparation) at a 2:1 (fungi:cell) ratio, followed by incubation at 37 ºC, 5 percent CO2 for 1 hour. Cells were then analyzed using a Cytoflex LX flow cytometer (Beckman). Percent phagocytosis was determined by dividing the total number of AF633+ HL-60 cells by the total number of HL-60 cells in each sample.

### Immunoblot Analysis

For each sample, 106 cells were lysed using cold buffer containing 50 mM Tris-HCl, 0.15 M NaCl, 1 mM EDTA, 1 percent NP-40, and 0.25 percent sodium deoxycholate. Gβ4 expression was assessed using monoclonal antibodies against the human ortholog (Abcam, ab223113), with GAPDH (clone D16H11, Cell Signaling Technology) serving as the loading control. IRDye 800 anti-mouse and IRDye 680 anti-rabbit secondary antibodies were used for visualization on Li-Cor Odyssey LX machine.

### Micropipette migration assay

Differentiated ΔGβ4 and WT HL-60 cells were stained with Hoescht (MilliporeSigma) for 20 min at 25 ºC and allowed to attach to fibronectin-coated 35 mm glass-bottom cover slips for 1 h (250,000 cells per dish). Samples were washed to remove nonadherent cells and left in 3 mL of complete RPMI media prior to imaging. A femptotip (Eppendorf) micropipette loaded with sterile filtered RPMI containing 2 percent fatty acid free BSA, 1 percent Pen/Strep, 200 nM fMLF, and trace AF647 dye was attached to a FemptoJet 4i (Eppendorf), and the micropipette tip positioned and the center of the imaging frame. Pressure (400 psi) was then applied to dispense the chemoattractant mixture and serial images (40 s intervals between frames) collected over the course of an hour at 20× magnification using a Ti Eclipse microscope with a CSU-0W1 Yokogawa camera (Nikon) at 37 ºC, 5 percent CO2. To measure migration after MP uptake, ΔGβ4 and WT HL-60 cells were preincubated with pHrodoGreen or pHrodoRed labeled MPs, respectively, for 3 hours. Cells that had taken up MPs during this time were FACS sorted based on green (ΔGβ4) or red (WT) fluorescence. A 1:1 mixture of these cells was then applied to fibronectin coated cover slips (250,000 cells per dish) and chemotaxis toward a point source of fMLF measured as described above.

### Scanning electron microscopy (SEM)

HL-60 cells adhered to fibronectin coated 1.5 coverslips (Fisher Scientific) were prepared for SEM with critical point drying. Briefly, samples were dehydrated with ethanol gradient drying in five increments from 50 percent ethanol to 100 percent ethanol, for 10 minutes at each step. They were then subjected to critical point drying in a Bal-Tec CPD 030 device (Bal-Tec, USA) according to manufacturer instructions. Dried samples were sputter-coated using a Cressington 108 Auto Sputter Coater with a 5-10 nm thick layer of gold-palladium (Cressington Scientific Instruments, UK) and imaged using a Zeiss Sigma VP scanning electron microscope (Carl Zeiss, Germany) at 3 kV accelerating voltage, 5 mm working distance in In-Lens mode.

### FIB-SEM imaging and analysis

Cells were plated on Aclar coverslips coated with fibronectin and then fixed with 2 percent glutaraldehyde, 2 mM CaCl2 in 0.08 M sodium cacodylate buffer, pH 7.2. This primary fixation was followed by a ROTO protocol (Reduced Osmium-Thiocarbohydrazide-Osmium) as follows: cells were incubated for one hour in 1 percent OsO4, 1.25 percent potassium ferrocyanide in 0.1 M sodium cacodylate buffer on ice, washed with buffer, and then incubated with 1 percent thiocarbohydrazide in water for 12 minutes. After washing, cells were treated with 1 percent Osmium tetroxide in 0.1 M cacodylate buffer for 30 min on ice. The samples were then dehydrated using a graded series of ethanol solutions and embedded in Eponate 12. Sample blocks were trimmed and then mounted on an SEM sample holder using double-sided carbon tape (EMS). Colloidal silver paint (EMS) was used to electrically ground the sides of the resin block. The entire surface of the specimen was then sputter coated with a thin layer (5 nm) of gold/palladium. The sample was imaged using immersion, TLD, back-scattered electron (BSE) mode on an FEI Helios Nanolab 650 microscope. Images were recorded after each round of ion beam milling using the SEM beam at 2.0 keV and 0.10 nA current with a working distance of <5 mm. Ion beam was held at 30 keV, with a milling current of 80 pA. Data acquisition occurred through automation using Auto Slice and View G3 software. Raw images were 4,096 × 2,048 pixels, with 20-50 nm slices viewed at a -38° cross-sectional angle. Each raw image had a horizontal field width (HFW) of 10-15 μm with an XY pixel size of 2-4 nm and 40 nm Z-step size. Images were aligned using the image processing programs in IMOD. All segmentation was performed using semi-manual thresholding and manual annotation of a test set (10 percent of data) using the LabKit machine learning plug-in for Imaris. 3-D renderings, reconstructions, surface/volume calculations, and internal vesicle volume calculations were performed using Imaris.

### Myriocin sphingolipid synthesis inhibition assays

For acute myriocin treatment, HL-60 cells on the 5th day of DMSO differentiation were treated with a final concentration of 50 nM myriocin (Millipore Sigma Aldrich) for 12 h prior to phagocytosis or C-trap experiments. For long term inhibition of sphingolipid synthesis, HL-60 progenitor cells were incubated over the 5 day DMSO differentiation in the presence of 50 nM myriocin. Treated cells were then used in phagocytosis or C-trap experiments. No observable effect on viability was observed in either treatment regime.

### Osmotic shock experiments

HL-60 cells or terminally differentiated primary human macrophages (derived from hiPSC cells) were suspended in 200 μL complete RPMI (10 percent FBS, 1.5 percent DMSO, 1 percent Pen/Strep) or RPMI 1650 (h-mCSF, 10 percent FBS, 1 percent Pen/Strep), respectively, and then applied to a Vi Cell Blue (Beckman Coulter, MA) cell counter to measure cell diameter. The samples were then osmotically shocked for 2 min or 4 min respectively by adding 180 μL of sterile ddH2O to 20 μL of suspended cells (final concentration of 2 × 105 cells per mL) before loading onto the same device. Each measurement was performed using 9 biological replicates. All recorded samples had >90 percent viability during the assay, as monitored by trypan blue (ThermoFisher) exclusion.

### Total internal reflection fluorescence (TIRF) imaging of frustrated phagocytosis

The inner 4 chambers of an 8 well chamber slide (Ibidi) were coated with biotinylated poly-L lysine, followed by streptavidin and biotinylated human IgG (Jackson ImmunoResearch 015-060-003) using an established protocol(Quann et al. 2009). Coated chamber slides were mounted on an Elyra 7 microscope (Zeiss) equipped with a 60× TIRF objective lens, followed by addition of ΔGβ4 or wild type HL-60 cells expressing F-tractin-mCherry and stained with CFSE (ThermoFisher Scientific). Samples were imaged in TIRF mode for 10 min using 15 s intervals at 37 ºC, 5

### Generation of mutant hiPSC lines

The 731.2B-iPSC line was generated from human fibroblasts (GM00731) purchased from the Coriell Institute(Miller et al. 2013). These cells have been stocked and characterized by the Stem Cell Research Facility at MSKCC. They are mycoplasma free, express the expected levels of OCT4, SOX2, and NANOG, and have a normal karyotype. GNB4-/- iPSCs were generated by CRISPR-Cas9 targeting. An sgRNA (TGCGAACAAGACGTACACTG) specific for GNB4 was first subcloned into the pSpCas9(BB)-2A-Puro (PX459) V2.0 vector (Addgene plasmid 62988). 731.2B-iPSC cells were then dissociated using Accutase (Innovative Cell Technologies) and electroporated with the sgRNA-Cas9 plasmid using an Amaxa 4D-Nucleofector (Lonza), following manufacturer’s instructions. Electroporated cells were cultured in 1 mL Essential 8 media (ThermoFisher Scientific) and Stemflex media (Stem cell technologies) at 37 ºC, 5 percent CO2, for 4 days and then dissociated and subcloned. Approximately 10 days later, individual colonies derived from single cells were picked, mechanically disaggregated and replated into two individual wells of a 96-well plate. Colonies were screened by PCR for biallelic frameshift mutations at the targeting site. Wild type clonal lines from the same targeting experiments were included as controls.

### hiPSC embryoid body formation and human primary macrophage differentiation

Macrophages were differentiated from hiPSCs using an established protocol(Lachmann et al. 2015), with modifications. Briefly, hiPSCs were seeded on CF1 mouse embryonic feeders and maintained for 3 days in ESC medium (KO-DMEM (10829-018), 20 percent KO-Serum (10828-028), 2mM L-glutamine (25030-024), 1 percent Nonessential Amino Acids (11140-035), 0.2 percent beta-mercaptoethanol (31350-010), 1 percent Pen/Strep (15140163) (all supplied by ThermoFisher) with 10 ng/mL bFGF (Peprotech 100-18B) and subsequently for 4 days in ESC media without bFGF. 7 days after seeding, at day (D)0 of the differentiation process, hiPSC colonies were detached in clusters using a 13 minute incubation with collagenase type IV (250 IU/mL final concentration) (Thermo Fisher Scientific; 17104019) and transferred to 6 well low adhesion plates in ESC media supplemented with 10 μM ROCK Inhibitor (Sigma; Y0503) to initiate the differentiation process. The plates were kept on an orbital shaker at 100 rpm for 6 days to allow for spontaneous formation of embryoid bodies (EB) with hematopoietic potential. At D6 of the differentiation process, well-formed 200-500 μm EBs were picked under a dissecting microscope and transferred onto adherent tissue culture plates (2.5 EBs/cm2) for cultivation in Hematopoietic Differentiation (HD) medium (APEL2 (Stem Cell Tech 05270), Protein free hybridoma medium (ThermoFisher Scientific; 12040077), 1 percent pen/strep, 25 ng/mL hIL-3 (Peprotech; 200-03), and 50 ng/mL hM-CSF (Peprotech, 300-25)). Starting from D18 of the differentiation and then every week onwards for up to two months, macrophages produced by EBs were carefully collected from suspension, filtered through a 100 μm mesh, plated at a density of 10,000 cells/cm2, and cultivated for 6 days in RPMI1640/GlutaMax (ThermoFisher Scientific; 61870036) medium supplemented with 10 percent FBS (EMD Millipore TMS-013-B), and 100 ng/mL human M-CSF before use in downstream experiments. All cells were cultured at 37 ºC, 5

### Membrane tension measurements

Plasma membrane tension was quantified using a C-Trap optical trapping device (Lumicks BV, Netherlands). An IR laser beam (50 mW, 1064 nm) was tightly focused through a series of mirrors, beam expanders and a high numerical aperture objective lens (63×/1.2 NA, Nikon Instruments) to form a steerable optical trap. Cells were immobilized inside an Ibidi μ-slide (Ibidi GmbH, Germany) treated with 200 μg/mL fibronectin (Thermo/Sigma). To measure plasma membrane tension, polystyrene beads (2.2 μm, Spherotech Inc, IL) were coated with concanavalin A (50 μg/mL, Thermo/Sigma) and added to the cell culture medium inside the slide. Beads were momentarily placed in contact with the cell membrane, and tethers were then extruded by moving the bead away from the cell perpendicularly at a speed of 2 μm/s. Force measurements were made using the Lumicks Bluelake software suite by capturing the exiting trapping light with a high numerical aperture condenser lens (63×/1.45, oil immersion, Zeiss AB, Germany) and measuring bead displacement in the trap with position-sensitive detectors through back focal plane interferometry. Membrane tether breaking was documented as a sharp discontinuity in tether force during tether extrusion, with breaking distance measured from simultaneously collected brightfield images using FIJI. Data analysis was performed using Python 3.8.0.

### RNA-sequencing

Cell samples were lysed in TRIzol (ThermoFisher) and the RNA extracted using the MagMAX mirVana Total RNA Isolation Kit (ThermoFisher catalog A27828) on a KingFisher Flex Magnetic Particle Processor (Thermo Scientific catalog 5400630) according to the manufacturer’s protocol. After RiboGreen quantification and quality control using an Agilent BioAnalyzer, 2 ng total RNA with RNA integrity numbers ranging from 8.5 to 10 was amplified using the SMART-Seq v4 Ultra Low Input RNA Kit (Clonetech catalog 63488), with 12 cycles of amplification. Subsequently, 5 13194*B*)*or*10((12227*B*,13028*B*) *ngofamplifiedcDNAwasusedtopreparelibrarieswiththeKAPAHyperPrepKit*(*KapaBiosystemsKK*8504 *seqdatawasconductedinRv*.4.0.5, *andcountnormalizationanddifferentialgeneexpressionwasperformedusingtheRpackageDESeq2.Atleast3independentreplicatesw*

### Lipidomics

Frozen cell pellets were thawed and extracted using a modified Folch protocol(Folch, Lees, and Sloane Stanley 1957). In brief, samples were resuspended in 300 μL of methanol containing SPLASH LIPIDOMIX (Avanti Polar Lipids) as internal standards, vortexed, and then mixed with 600 μL of chloroform. 180 μL of water was added to each tube to induce phase separation, and after mixing, samples were centrifuged at 16,000 × g for 5 min at 4 ºC. Subsequently, the lower, chloroform layer was collected and the aqueous layer re-extracted using 450 μL of chloroform:methanol:water (3:48:47 v/v/v). The lower, chloroform layer was collected and pooled with the previous extract. Samples were then dried under nitrogen at 40 ºC and resuspended in 100 μL of 90:10 methanol:chloroform. Lipid profiling was performed using an Agilent 6546 Q-TOF mass spectrometer in positive and negative ionization modes, coupled to a ZORBAX Eclipse Plus C-18 Column (100 mm × 2.1, 1.8 μm particle size, Agilent). Mobile Phase A consisted of 10 mM ammonium formate in 50:30:20 water:acetonitrile:isopropanol.

Mobile Phase B consisted of 10 mM ammonium formate in 1:9:90 water:acetonitrile:isopropanol. LC gradient conditions were: 0 min at 0 percent B; 2.7 min at 45 percent B; 2.8 min at 53 percent B; 9 min at 65 percent B; 9.10 min at 89 percent B; 11 min at 92 percent B; 11.10 min 100 percent B; 12 min at 10 percent B; 15 min at 10 percent B. Other LC parameters were: flow rate at 0.4 mL/min, column temperature at 60 ºC and injection volume was 5 μL. MS source parameters included: gas temp: 200 ºC; gas flow: 10 L/min; nebulizer pressure: 50 psig; sheath gas temp: 300 ºC; sheath gas flow: 12 L/min; VCap: 3000 V; nozzle voltage: 0 V; fragmentor: 150V. For lipid annotation, 5 injections of iterative MS/MS acquisition were performed on a pooled lipid extract in both positive and negative polarity. To identify lipid species, iterative MS/MS acquisition data in positive and negative polarity was processed using Agilent Lipid Annotator software 1.0(Koelmel et al. 2020). Subsequent targeted feature extraction and peak integration was performed using Skyline(Adams et al. 2020). For statistical analysis, the Welch T-test was used for pairwise comparisons between wild type and Gβ4 groups with Benjamini-Hochberg FDR correction.

### Generation of Gnb4-/- mice

The animal protocols used for this study were approved by the Institutional Animal Care and Use Committee of MSKCC. Two sgRNA sequences (5’-cgtcaaaatatcgcaagtgc-3’) and (5’-aggtgtcagatcaaacc-3’) targeting exon 4 of the Gnb4 locus (IDT) were injected together with purified Cas9 protein (IDT) into C57BL/6 zygotes, which were then transferred into C57BL/6 pseudopregnant females. Founder animals were screened for deletion of the entire exon 4 and the presence of an early stop codon in exon 5, and then bred to homozygosity. Routine genotyping PCRs were performed using the following forward (5’-ggagaacagctagtactcttaac-3’) and reverse (5’-aaaagtatttattagcagtatc-3’) primers. The resulting amplicons for WT and Gnb4 KO mutant alleles are 1360 bp and 305 bp respectively.

### a. fumigatus FLARE intratracheal mouse fungal infections

Gnb4-/- and WT mice were infected by intratracheal (i.t.) administration of 60 × 106 FLARE Af293 A. fumigatus (see procedure above for staining protocol) in 50 μL. WT mice were also infected with unlabeled A. fumigatus to serve as analysis controls (see below). 18 h post infection, mice were euthanized and their lungs harvested into 5 mL of digestion buffer (PBS with 5 percent FBS, 0.1 mg/mL DNAse I, and Type IV collagenase (Worthington) at 2.2 mg/mL). Tissue homogenization was performed using a MACSTm Octo Dissociator for 55 sec at 1302 rpm, followed by slow rotation at 37 ºC for 40 min. A final mechanical homogenization was performed for 37 sec at 2079 rpm, after which all samples were diluted with 5 mL of PBS + 5 percent FBS, filtered through a 100 μm pore size cell strainer, and centrifuged at 300 × g for 5 min at 4 ºC. The pellet was resuspended in 2 mL of ACK lysis buffer (BD) and incubated for 15 min to achieve red blood cell lysis. After quenching in 4 mL of RPMI-1640 +10 percent FBS, cells were centrifuged at 300 × g for 5 min at 4 ºC, followed by resuspension in cold FACS buffer (PBS + 5 percent FBS). To assess dissemination at 24 h, lungs, dLN, spleen, and 200 μL of blood were harvested from each mouse. Lungs were processed as described above. dLN and spleen were homogenized and filtered through a 100 μm pore size cell strainer. Subsequently, the spleen single cell suspension was centrifuged and resuspended in 2 mL of ACK lysis buffer (BD) and incubated for 15 min to achieve red blood cell lysis. After quenching in 4 mL of RPMI-1640 +10 percent FBS, cells were centrifuged at 300 × g for 5 min at 4 ºC, followed by resuspension in cold FACS buffer (PBS + 5 percent FBS). Blood samples were diluted into 2 mL of ACK lysis buffer (BD) containing 100 mM EDTA and incubated for 15 min to achieve red blood cell lysis. After quenching in 4 mL of RPMI-1640 +10 percent FBS, cells were centrifuged at 300 × g for 5 min at 4 ºC, followed by resuspension in cold FACS buffer (PBS + 5 percent FBS). Aliquots of 2 × 106 cells from each organ were dispensed into round bottom 96 well plates (Corning) and then stained for 20 min at 4 ºC with a viability dye (Tonbo Ghostdye Violet 510 1:250) along with antibodies against Ly6G (BUV395, 1:100), CD11b (BUV805, 1:100), Siglec F (BV650, 1:100), CD45 (BV785), and CD11c (PE-Cy7, 1:100). Samples were then applied to a CytoFlex LX flow cytometer and analyzed using FlowJo software. Neutrophils were identified as CD11b+Ly6G+SiglecF-CD11c-cells. Percent phagocytosis was determined by dividing the total number of AF633+ neutrophils in each sample by the total number of neutrophils. Percent FLARE killing was expressed as the number of AF633+dsRed-neutrophils over the total number of AF633+ neutrophils in each sample. Gating for AF633+ neutrophils was facilitated using unlabeled neutrophils extracted from mice infected with unlabeled A. fumigatus. To quantify fungal infection, lung suspensions were diluted 50-fold and 50 μL of each sample was plated on Sabouraud dextrose agar (ThermoFisher). CFUs were quantified after 2 d of incubation at 37 ºC. Fluorochromes Antigen Company Clone number dilution BUV395 Ly6G BD Biosciences 1A8 1:100 BUV805 CD11b BD Biosciences M1/70 1:100 BV510 L/D Aqua Tonbo Ghostdye Violet 510 1:100 BV650 SiglecF BD Biosciences E50-2440 1:100 BV785 CD45 BioLegend 30-F11 1:100 PE-Cy7 CD11c BioLegend N418 1:100

### Statistics

Graphpad was used for statistical analysis (Graphpad software, Inc). Details can be found in the legend of each figure. N represents number of independent biological replicates.

